# The locomotor behavior of subfossil Malagasy sloth-lemurs (Strepsirrhini: Palaeopropithecidae) and koala-lemurs (Strepsirrhini: Megaladapidae): new insights from limb trabecular bone

**DOI:** 10.1101/2025.10.21.683631

**Authors:** Fabio Alfieri, Julia Arias-Martorell, Carla Argilés-Esturgó, Damiano Marchi

**Author notes:** **Corresponding author**: Fabio Alfieri,;, address: Institute of Ecology and Evolution, Universität Bern, Baltzerstrasse 6, 3012, Bern, Switzerland.

## Abstract

The locomotion of Malagasy Quaternary subfossil lemurs—including palaeopropithecids (‘sloth-lemurs’) and megaladapids (‘koala-lemurs’)—has been investigated on abundant postcranial remains. Proposed strategies include some without living primate parallels, including sloth-like suspensory arboreality in palaeopropithecids, although the degree of suspensory behaviour in palaeopropithecids, or locomotor diversity in koala-lemurs are poorly understood. Differently from external morphology, internal bone structure in these taxa is largely unexplored. We compared the humeral and femoral trabecular architecture of sloth- and koala-lemurs to several extant mammals, studying spherical trabecular samples extracted from high-resolution scans. After defining locomotor categories from quantitative data, we tested links between trabecular parameters and locomotor modes through exploratory and multivariate analyses, accounting for body size and phylogeny. In extant mammals, only femoral trabecular traits—particularly Degree of Anisotropy and Bone Volume Fraction—were significantly associated with locomotion, distinguishing suspensory and bridging arboreal taxa from others. Using this model, we inferred suspensory adaptations in palaeopropithecids, especially *Palaeopropithecus*—confirming earlier reconstructions—but also in *Megaladapis edwarsi*, a striking result that would make *M. edwarsi* the largest mammal ever known to adopt such habits, approached only by extant orangutans. This work highlights the potential of internal bone structure for reconstructing primate locomotor evolution.

## Introduction

Studying the fauna of Madagascar offers a unique opportunity to address central questions in biology, due to the island’s exceptional biodiversity and distinctive geological and evolutionary history (e.g. Crowley, 2010; Muldoon, 2010; Muldoon *et al*. 2009). Lemurs, among the most iconic Malagasy organisms, are taxonomically diverse (representing ∼13% of living primates, Martin, 2000; Rowe, 1996) and ecologically varied (Martin, 2000), particularly in their locomotor/postural behaviors. Among others, these include climbing and leaping, quadrupedalism (arboreal and terrestrial), vertical clinging, and terrestrial hopping (Demes *et al*. 1996; Furnell *et al*. 2015; Gebo, 1987; Jungers *et al*. 2005; Walker, 1979, 1974).

When humans first reached Madagascar (∼2300 years ago; Muldoon, 2010, see also Douglass *et al*. 2019), they encountered not only the modern primate community but also additional nowadays extinct lemur groups—i.e. palaeopropithecids (“sloth-lemurs”), archaeolemurids (“monkey-lemurs”), megaladapids (“koala-lemurs”), the daubentoniid *Daubentonia robusta*, and the lemurid *Pachylemur*. These species persisted for centuries alongside humans, with some surviving until ∼500 years ago (Crowley, 2010). Their recent extinction led to the preservation of extensive skeletal material which has been pivotal in reconstructing their biology, including body mass (e.g. Jungers *et al*. 2002, 2008), phylogenetic affinity (e.g. Herrera and Dávalos, 2016; Karanth *et al*. 2005; Kistler *et al*. 2015; Yoder *et al*. 1999) and locomotor/positional behaviors (e.g. Godfrey *et al*. 1995; Jungers *et al*. 1997; Shapiro *et al*. 2005, see also below). The latter resulted remarkably diverse, as shown by postcranial material, and, if combined with the locomotor diversity observed in extant lemurs, the whole lemur behavioral range approaches that of all other primates together (Godfrey *et al*. 1997; Granatosky, 2022; Jungers *et al*. 2002, 2005; Tattersall, 1982; Walker, 1974).

Notably, within subfossil lemurs, sloth-lemurs and koala-lemurs share inferred ecological and physiological traits—i.e. slow arboreality, low metabolic rates, and dominance of rest in daily activity (Alfieri *et al*. 2021, 2023; Godfrey *et al*. 2016; Hogg *et al*. 2015; Walker *et al*. 2008). For instance, the radius of curvature of their semicircular canals—a proxy for locomotor agility (Spoor *et al*. 2007) indicates they were the least agile among subfossil lemurs (Walker *et al*. 2008). Such ecological traits, shared with tree-sloths and koalas, may be associated with convergent morphologies such as low cortical bone compactness in limb diaphyses (Alfieri *et al*. 2021). However, other anatomical features suggest biomechanical divergence between sloth-lemurs and koala-lemurs, despite their broad ecological similarities (Alfieri, 2022; Alfieri *et al*. 2023).

Sloth-lemurs—*Palaeopropithecus*, *Mesopropithecus*, *Babakotia*, and *Archaeoindris* (Godfrey and Jungers, 2003; Gommery *et al*. 2009; Jungers *et al*. 1997; Simons *et al*. 1992)—ranged from ∼11 kg to ∼160 kg of body mass (Granatosky, 2022); indriids are their closest living relatives (Herrera and Dávalos, 2016; Karanth *et al*. 2005; Kistler *et al*. 2015; Tattersall, 1973; Tattersall and Schwartz, 1974). While the locomotion of *Archaeoindris* remains debated due to limited fossil evidence (e.g. Godfrey *et al*. 2016; Godfrey and Jungers, 2002), abundant studies of postcranial axial and appendicular skeletal traits have suggested that *Palaeopropithecus*, *Mesopropithecus*, *Babakotia* likely showed specialized suspensory adaptations based on below-branch quadrupedalism, and that this specialization was potentially more extreme in *Palaeopropithecus*, followed by *Babakotia*, which would have been intermediate, and finally by the least-specialized *Mesopropithecus* (Godfrey *et al*. 1995; Godfrey and Jungers, 2003; Granatosky, 2018, 2020, 2022; Hamrick *et al*. 2000; Jungers, 1980; Jungers *et al*. 1991; Marchi *et al*. 2016; Patel *et al*. 2013; Shapiro *et al*. 1994, 2005; Wunderlich *et al*. 1996). While there seems to be a certain consensus on the more extreme adaptations of *Palaeopropithecus*, the relative degree of specialization in suspensory behaviors of the other two taxa appears less clear. For instance, it was also proposed that *Babakotia* and *Mesopropithecus* —the smaller palaeopropithecids—were more similar to the two-toed sloth *Choloepus*, contrasting with *Palaeopropithecus* that was instead inferred to resemble more the three-toed sloth *Bradypus* (Marchi *et al*. 2016 and references). Also, *Babakotia* and *Mesopropithecus* are not distinct for some potentially functional traits (i.e., within-limb proportions), hence they do not clearly reflect the expected different degree of suspensory adaptations mentioned above (Marchi *et al*. 2016). Accordingly, *Babakotia* and *Mesopropithecus* have been overall described as taxa with a more diverse locomotor repertoire, compared to *Palaeopropithecus* (Granatosky, 2022).

Koala-lemurs, i.e. *Megaladapis* spp. (Gebo, 1986; Jungers, 1977, 1980; Szalay and Delson, 1979, 1979; Tattersall, 1975; Walker, 1967), include three species with body mass ranging from the ∼45 kg of *M. madagascariensis* to the ∼85 kg of *M. edwarsi*, with the medium-sized *M. grandidieri* (∼75 kg) in between (Jungers, 1978, 1977; Jungers *et al*. 2008; Tattersall, 1975; A. Walker, 1967). The phylogenetic position of *Megaladapis* spp. has been controversial—i.e., close to *Lepilemur* (Godfrey *et al*. 2010; Tattersall and Schwartz, 1974), to Lemuridae (Karanth *et al*. 2005; Kistler *et al*. 2015; Orlando *et al*. 2008), or early diverging from all non-daubentoniid lemurs (Herrera and Dávalos, 2016). Here, we will consider koala-lemurs as the sister taxon of Lemuridae, since this has been confirmed by more recent works addressing nuclear DNA sequences (Marciniak *et al*. 2021; Upham *et al*. 2019). The locomotor behavior of *Megaladapis* spp. was long debated, too (Godfrey and Jungers, 2002). While there is now wide consensus on their arboreality with behaviors dominated by forelimb activity (but without being characterized by extreme suspensory adaptations such as in palaeopropithecids, see above) — which is supported by a high intermembral index, powerful hands and feet, and body proportions reminiscent of koalas (Godfrey *et al*. 2016; Godfrey and Jungers, 2002; Jungers, 1978; Wunderlich *et al*. 1996) — some uncertainties remain regarding their specific postural and locomotor behaviors. A first traditional view holds that vertical climbing and clinging were the primary postures for all the *Megaladapis* taxa (Jungers, 1977, 1978; agreeing with Carleton, 1936). In this framework, the three species are merely considered as size and geographic variants— i.e., the largest *M. edwarsi* and the smallest *M. madagascariensis* representing the south-southwestern varieties, with the medium sized *M. grandidieri* living in central Madagascar (Jungers, 1978, 1977; Jungers *et al*. 2008; Tattersall, 1975; A. Walker, 1967). However, new fossil evidence has led to the recognition of two potentially behaviorally distinct sub-lineages of koala lemurs: the small to medium sized and widely distributed *Megaladapis* (*M. madagascariensis* + *M. grandidieri*)—characterized by vertical climbing combined with quadrupedal hanging and pedal suspension (Jungers *et al*. 2008, 2002b; Wunderlich *et al*. 1996, 1994)—and the large-sized and southern *Peloriadapis* (sub-genus corresponding to *M. edwarsi*; Vuillame-Randriamanantena *et al*. 1992; Wunderlich *et al*. 1996), proposed as relatively more terrestrial compared to *M. madagascariensis* and *M. grandidieri* (Jungers *et al*. 2008, 2002b; Wunderlich *et al*. 1994,1996).

Most of the reconstructions of locomotion in subfossil lemurs have focused on bone external morphology (e.g., Godfrey *et al*. 1995; Marchi *et al*. 2016; Wunderlich *et al*. 1996; although Marchi *et al*. analyzed diaphyseal cross-sectional properties too). However, outer morphology, especially concerning joint anatomy, is potentially more genetically (thus, less functionally) driven than internal bone structure, i.e. the inner repartition of osseous tissue within skeletal elements (Lieberman *et al*. 2001; Rafferty and Ruff, 1994). It is especially relevant in the long bones of the appendicular skeleton—known for their potential to inform on locomotor behavior (Dunn, 2018). Indeed, while long bone external morphology is more representative of the potential niche—i.e. what organisms can theoretically do in terms of locomotor habits (e.g. the maximum range of rotation allowed at a joint without causing disarticulation or injury)—inner structure more closely reflects the realized niche—i.e. what organisms actually do and which behaviors are commonly performed by a species. This applies to all levels of inner architecture, such as diaphyseal compact bone and epiphyseal trabecular structure. Particularly the latter has been shown (both experimentally, e.g., Biewener *et al*. 1996; Pontzer *et al*. 2006; and computationally, e.g., Huiskes *et al*. 2000; Keaveny *et al*. 2001) to adapt in response to biomechanical loadings (see also Kivell, 2016 and references therein). This characteristic highlights the potential of the trabecular structure of limb long bones to reflect the loading conditions to which they are subjected, which can, in turn, be linked to specific locomotor behaviors. Some studies have not found trabecular structural differences reflecting different locomotor habits (e.g., Carlson *et al*. 2008; Ryan and Walker, 2010), which was explained by the fact that trabecular bone might potentially also affected by non-functional aspects (e.g. systemic; Tsegai *et al*. 2018, genetic, Ryan *et al*. 2017; ontogenetic; Macho *et al*. 2005; and hormonal; Khosla *et al*. 2006). However, many comparative analyses across several postcranial regions have shown that trabecular bone co-varies with locomotor behavior (e.g., Alfieri, 2022; Alfieri *et al*. 2022; Arias-Martorell *et al*. 2021; Chang *et al*. 2008; Fajardo and Müller, 2001; Georgiou *et al*. 2019; Mielke *et al*. 2018; Ryan and Shaw, 2012; Saparin *et al*. 2011; Tsegai *et al*. 2013), making the study of this anatomical level a common approach for inferring past locomotor habits in extinct primates (e.g. Barak *et al*. 2013; Kivell *et al*. 2018; Rook *et al*. 1999; Ryan *et al*. 2018; Ryan and Ketcham, 2002a). Despite its potential, the internal architecture of the long bones of the sloth-lemurs and koala-lemurs has not yet been analyzed. Only recently, the trabecular structure of long bone epiphyses in palaeopropithecids and *Megaladapis* has been quantified, but primarily to investigate evolutionary patterns in slow arboreal mammals (Alfieri *et al*. 2023), rather than to directly infer their locomotor behavior.

To fill this gap, in this study we examine humeral and femoral trabecular architecture in sloth-lemurs and koala-lemurs, comparing them with those of a sample of extant mammals encompassing: potential ecological analogues of subfossil lemurs, inferred sister taxa of sloth-lemurs and koala-lemurs and other relevant groups. We first used quantitative behavioral data on extant taxa to classify them in locomotor categories; we then analyzed their humeral and femoral trabecular structure with the aim of understanding which skeletal region and trabecular trait discriminate extant mammals according to their locomotion. Hence, we inferred the most likely locomotor behavior for extinct lemurs, using both an exploratory—i.e., assessing morphospaces—and analytical approach—i.e., running a phylogenetically informed discriminant function analysis on the most informative trabecular traits. This protocol allowed us to understand whether trabecular bone supports previous hypotheses regarding: 1) the particularly derived suspensory adaptations of *Palaeopropithecus*; 2) the relatively intermediate degree of suspensory specialization in *Mesopropithecus* and *Babakotia*; and, 3) potential locomotor differences across the small/medium sized *M. madagascariensis* and *M. grandidieri*, and the large sized *M. edwardsi* (see above).

## Materials and Methods

### Sample

#### Subfossil lemurs

We used high-resolution μCT data derived from humeri and femora of Malagasy subfossil lemurs collected in the Muséum national d’Histoire Naturelle (Paris, France) and the Division of Fossil Primates, Duke Lemur Center, Durham (NC, USA). Details regarding acquisition can be found in Alfieri *et al*. (2023). The studied data derive from specimens belonging to *Babakotia* sp. (one complete humerus and one distal humerus), *Palaeopropithecus* sp. (four complete humeri, one proximal humerus, one distal humerus, four complete femora, one proximal femur, two distal femora), *Mesopropithecus dolichobrachion* (one complete humerus), and *Megaladapis* sp. (three complete humeri, one proximal humerus, one distal humerus, two complete femora, two proximal femora and two distal femora). *Babakotia* sp. was assigned to *Babakotia radofilai*, the only species recognized within the genus, while *Palaeoproprithecus* sp. and *Megaladapis* sp. were assigned to known species within the two genera, following Alfieri *et al*. (2023) and as summarized in Supplementary Information (Appendix S1).

#### Extant primates

We built a comparative sample of extant mammal species collecting μCT data from previous studies (Alfieri *et al*. 2022, 2023; Arias-Martorell *et al*. 2021; Georgiou *et al*. 2018, 2019; Kivell *et al*. 2018; Ryan *et al*. 2018; Tsegai *et al*. 2018) and/or downloaded from MorphoSource (https://www.morphosource.org; Supporting Information Tables S1-S2). These data derive from 55 humeri and 53 femora collected across several institutions (Supporting Information, Appendix S2). We studied adult specimens, as determined by full epiphyseal fusion, and selected epiphyses with preserved trabecular bone for the extant sample. For the subfossil lemur epiphyseal data, perfect preservation of trabeculae was not always possible; therefore, we either discarded these specimens or adopted procedures to analyze fragmented individuals (see below). Most of the studied extant mammals have been proposed as extant ecological analogues of extinct lemurs and/or were used in previous morphological comparative works of the studied subfossil species (e.g. Alfieri *et al*. 2023, 2021; Amson and Nyakatura, 2018; Boyer *et al*. 2015; Godfrey, 1988; Godfrey *et al*. 1995, 2016; Granatosky, 2022; Granatosky *et al*. 2014; Hamrick *et al*. 2000; Jungers *et al*. 1997, 2002, 2008; Marchi *et al*. 2016; Shapiro *et al*. 2005; Walker *et al*. 2008; Wunderlich *et al*. 1996): i.e. tree-sloths (both three-toed, Bradypodidae, and two-toed sloths, Choloepodidae), the koala (Phascolarctidae), lorisids (Lorisidae) and non-human great apes (Hominidae). We also included representatives from the two families inferred as the extant closest relatives of palaeopropithecids and *Megaladapis*—i.e., Indriidae and Lemuridae, respectively (Baab *et al*. 2014; Herrera and Dávalos, 2016; Marciniak *et al*. 2021; Upham *et al*. 2019). Also, we included the platyrrhine *Alouatta caraya*, as an example of a non-strepsirrhine primate adapted to an energy-saving slow arboreal lifestyle (Bicca_Marques and Calegaro_Marques, 1998; Prates *et al*. 2018), an ecology that has been inferred for both palaeopropithecids and *Megaladapis* (Alfieri *et al*. 2021; Godfrey *et al*. 2016; Walker *et al*. 2008). Additonally, we included galagids, a strepsirrhine family that is not closely related to subfossil lemurs and whose members are specialized in vertical clinging and leaping (Fleagle, 2013; Gebo, 2011; Walker, 1979), the most common and possibly ancestral behavior in the strepsirrhine radiation (Boyer *et al*. 2017). By doing so, we were able to include a leaping group that is not closely related to extinct lemurs, contrary to indriids, which show leaping habits (see below) but are also the extant sister taxon of palaeopropithecids (Fig. 1). All taxa included in the sample were taxonomically categorized at the species level (more details in Tables S1-S2).

**Fig. 1.**
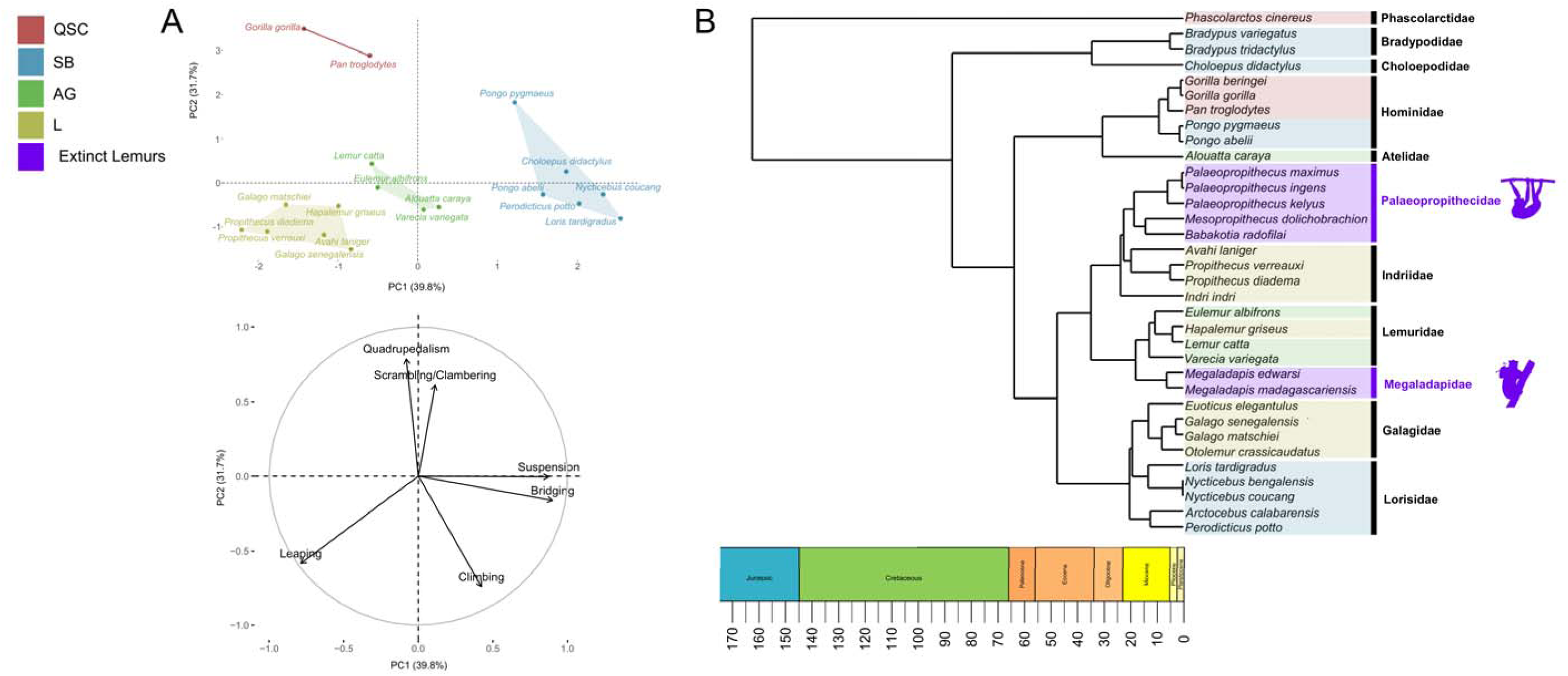
**A.** PCA on quantitative locomotor data (top: scatterplot; bottom: variable loading plot). Identified locomotor categories are: Leaping (L), Quadrupedal/Scrambling/Clambering (QSC), Suspensory/Bridging (SB) and Arboreal Generalists (AG). **B.** phylogenetic relationships between the studied species with locomotor categories mapped on the phylogeny.

#### Extant primates: locomotor diversity

For 62% of the extant species here included (Fig, 1), we were able to use data on the relative frequency (expressed in percentage) of locomotor behaviors collected by Granatosky (2018). In collecting these quantitative data, we largely followed the approach of Monclús-Gonzalo *et al*. (2023, 2025)—i.e., we pooled the behaviors quantified by Granatosky (2018) in six broader locomotor categories: 1) Leaping; 2) Quadrupedalism; 3) Climbing; 4) Suspension; 5) Bridging; 6) scrambling/clambering. The description of the locomotor categories can be found in Monclús-Gonzalo *et al*. (2023). We transformed the frequency data through the arcsine square root, a common procedure when proportional ecological data are studied (Monclús-Gonzalo *et al*. 2023; Sokal and Rohlf, 1995) (Table S3). For some species not listed by Granatosky (2018), we used quantitative data from closely related taxa (Appendix S3). Since we aimed to obtain quantitative data representative of species, in cases of multiple datasets for a single taxon reported by Granatosky (2018), we averaged the data for each behavior, resulting in one single value per locomotor category per species. The behavioral dataset was used to run a Principal Component Analysis (PCA, Table S4) from which we visually identified, on the PC1-PC2 biplot and variable loading plot, how taxa are divided into sub-groups due to their locomotor behaviors. These subgroups represented the discrete locomotor categories used to assign species for which quantitative locomotor data (*sensu* Granatosky 2018) were unavailable.

### Extraction of trabecular parameters

Most of the collected μCT data (i.e. all the species except for apes and *A. caraya*) were spherical Volumes of Interest (VOIs) of trabecular bone, that were extracted from head and capitulum of the humerus, and from head, lateral condyle and the medial condyle of the femur (Fig. 2) in previous works (Alfieri *et al*. 2022, 2023; as also summarized in Appendix S4). As for the rest of the sample, VOIs were extracted using the same protocols. Ultimately, all the studied VOIs represent homologous regions, i.e. the largest sphere including trabecular bone and not including cortical bone, extracted from the center of the humeral head, capitulum, femoral head, lateral and medial condyle (Fig. 2). The VOIs were imported in ImageJ2 (Rueden *et al*. 2017) and segmented, isolating the trabecular bone fraction. For extant species it generally corresponds to binarize the μCT data since only two fractions should be distinguished, i.e. trabecular bone from intertrabecular voids, to be assigned to foreground (i.e. 1) and background (i.e. 0), respectively. Hence, for almost all the extant species’ VOIs we used the automatic binarizing tool of ImageJ2 (‘Threshold’), yielding a result that we considered satisfying by assessing, through a visual comparison, the respective non-binarized VOIs. For the other VOIs (the few from extant species and those from subfossil lemurs), we followed alternative procedures, based on manual cleaning and/or tools for automated recognition of non-osseous fractions (detailed in Appendix S5). We then computed degree of anisotropy (DA), trabecular thickness (Tb.Th), average branch length (Av.Br.Len), bone volume fraction (i.e. BV/TV), bone surface density (i.e. BS/TV) and connectivity density (Conn. D). It was undertaken using BoneJ2 (Image plugin, Domander *et al*. 2021) and subsequent adjustments related to the use of spherical VOIs, as detailed in Appendix S6, where it is also detailed how we identified and discarded VOIs including a too low number of trabeculae. Given our research focus on subfossil lemurs, we aimed to maximize their representation in our sample. Therefore, we included some of their VOIs even if they contained peripheral regions with broken or missing trabeculae. To do so, we gradually reduced the size of these spherical VOIs until the biased regions were excluded. Notably, this approach requires applying the same scaling factor to all the other taxa to ensure consistency, i.e. studying anatomically homologous regions. However, reducing VOI sizes also decreases the number of trabeculae included. Thus, as justified in Appendix S7, for each epiphysis this procedure was balanced against the need to avoid underrepresenting some extant taxa due to the discarding deriving from too low trabecular number.

**Fig. 2.**
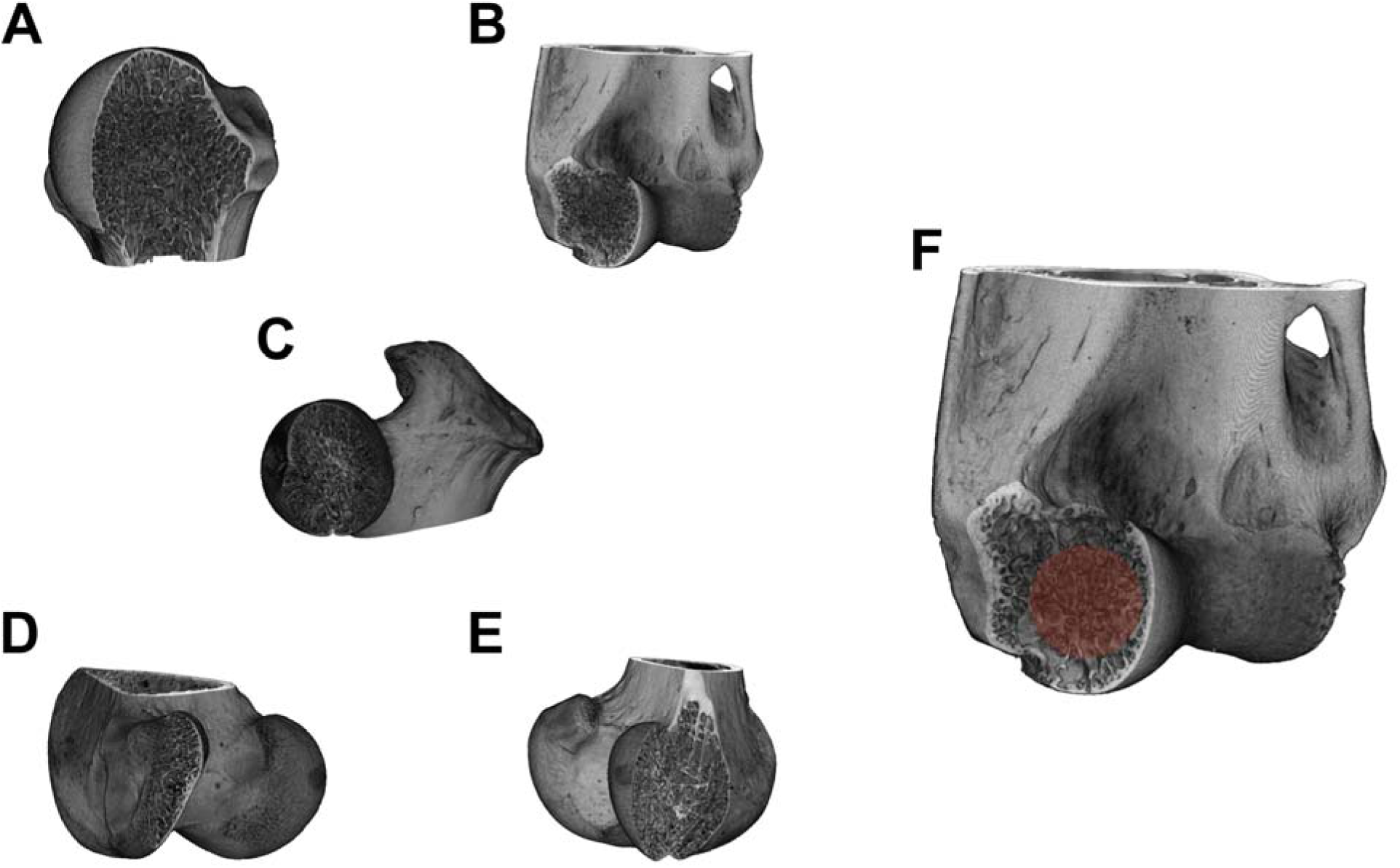
Humeral (**A.** proximal; **B.** distal) and femoral (**C.** proximal; **D.** distal lateral condyle; **E.** distal medial condyle) epiphyses from which trabecular bone was studied. **F.** The extraction of a spherical Volume of Interest (highlighted in orange) is exemplified. Images refer to the humerus of *Perodicticus potto* NMW 32674 and the femur of *Propithecus* sp AMNH 170474 and are not to scale.

### Statistical analyses

#### Relationship between trabecular structure and locomotion: inferential tests

Due to the high intercorrelation among trabecular parameters (e.g. Ryan and Shaw 2012), we analyzed them collectively as multivariate datasets, i.e. five anatomically separated datasets were defined: proximal humerus, distal humerus, proximal femur, lateral condyle, and medial condyle. We conducted a first set of PCAs on centered and scaled datasets including only trabecular parameters (’PCA_TP_’, hereafter). The resulting PC_TP_1s-PC_TP_4s (explaining the 93.4%-96.4% of the variance across the five datasets; Tables S5-S9) were used to test for a relationship between trabecular structure and ecological categories while accounting for phylogenetic affinities among species. As phylogeny, we used the time-calibrated tree deriving from the mammalian tree of Upham *et al*. 2019, adapted to our sample (see Appendix S8). We hence ran multivariate Phylogenetic Generalized Least Squares (mvPGLS) regressions (‘mvgls’ function, ‘mvMORPH’ package; Clavel *et al*. 2015) in R 4.3.1. (R Core Team, 2023) and phylogenetic MANCOVAs (‘manova.gls’ function, ‘mvMORPH’, Pillai’s trace tests) (Table 1). Beyond locomotion and phylogeny, trabecular bone is also potentially affected by allometry. To account for this source of variation too, we included a body mass proxy (BMp) as co-variate in mvPGLSs. As detailed in Appendix S9, as BMp we took the centroid size, deriving from the landmark coordinates from Alfieri *et al*. 2022, 2023, predicted from a linear regression of BMp against bone metric measurements for specimens lacking centroid size values. To establish whether in mvPGLSs we needed to account for ecology-BMp interactions too, we preliminarily tested for a relationship between BMp and locomotor categories (phylogenetic ANOVA, ‘phylANOVA’ function, ‘phytools’ package, Revell, 2012)

**Table 1.** Results from mvPGLSs and pMANCOVAs testing the relationship between trabecular structure (quantified through PC_TP_1-PC_TP_4 at different anatomical regions) and locomotion, while accounting for body mass effects (see Methods for details)

| **Anatomical region (articulation)** | **Principal Components** | **Proportion of variance** | **Body mass *p*-value** | **Locomotion *p*-value** |
| --- | --- | --- | --- | --- |
| **Proximal humerus (humeral head)** | PC<sub>TP1</sub>-PC<sub>TP4</sub> | 93.4% | <0.001 | 0.21 |
| **Distal humerus (humeral head)** | PC<sub>TP1</sub>-PC<sub>TP4</sub> | 95.6% | <0.001 | 0.46 |
| **Proximal femur (femoral head)** | PC<sub>TP1</sub>-PC<sub>TP4</sub> | 95.4% | <0.001 | 0.094 |
| **Distal femur (lateral condyle)** | PC<sub>TP1</sub>-PC<sub>TP4</sub> | 96.8% | <0.001 | 0.024 |
| **Distal femur (medial condyle)** | PC<sub>TP1</sub>-PC<sub>TP4</sub> | 96.4% | <0.001 | 0.019 |

#### Relationship between trabecular structure and locomotion: qualitative assessment

We conducted an additional set of PCAs on centered and scaled datasets including trabecular parameters and BMp (’PCA_TP+BM_’, hereafter). BMp was included to visualize the effects of body size on trabecular structure and, by doing so, we could observe locomotor patterns on biplots built with PC_TP+BM_s that are not primarily driven by BMp. It could be detected by assessing how BMp contributes to PC_TP+BM_s in the respective variable loading plots (i.e. through the BMp vectors length and direction). Moreover, through this approach we avoided potential biases associated with univariate size correction (e.g., residuals), which may not fully remove size effects in multivariate analyses (e.g. Alfieri *et al*. 2025a). In PCA_TP+BM_s, we also included data of subfossil lemurs to discuss their position on scatterplots. We built biplots with the first three PC_TP+BM_s (Figs. 3-7, Tables S10-S14) and on them we observed the distribution of locomotion categories and which specific locomotor groups are discriminated by which specific trabecular properties. This included assessing whether subfossil lemurs occupy regions preferentially associated with certain extant taxa, thereby guiding our choice of relevant locomotor categories for further analysis (see Appendix S10). When we identified a trend related to a single PC_TP+BM_ (i.e. locomotor categories, or combinations of them, are mainly discriminated along one PC_TP+BM_), we tested for significance through univariate PGLS and ANOVA (‘gls’ function, ‘nlme’ package, Pinheiro *et al*. 2020). The contribution of trabecular traits and BMp to PC_TP+BM_s was assessed through eigenvector loadings and correlation analyses (Pearson coefficients and coefficients of determination, R^2^).

**Fig. 3.**
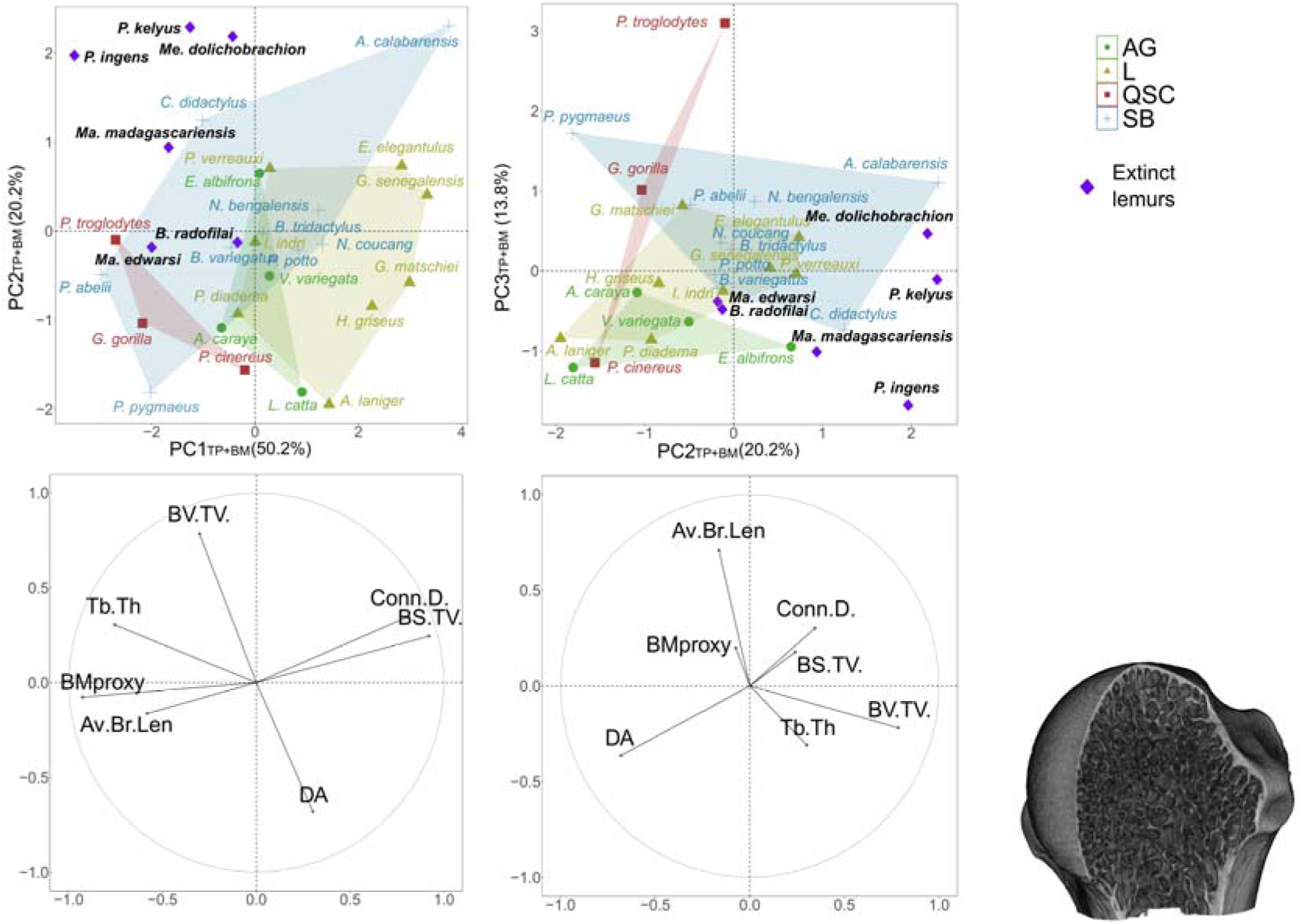
Proximal humerus PCA_TP+BM_ results. Top: scatterplots with extant species grouped by locomotor category and subfossil lemur data shown with purple rhombi. Bottom: corresponding variable loading plots.

#### Phylogenetic Discriminant Function Analysis

Once a subset of trabecular traits from different anatomical levels was identified as the most effective at discriminating specific locomotor categories (Table 2), we compiled a single heterogeneous dataset to run a phylogenetic Discriminant Function Analysis (pDFA). The pDFA preliminarily required a multivariate PGLS (‘mvgls’ function, ‘mvMORPH’) and phylogenetic MANOVA (‘manova.gls’ function, ‘mvMORPH’) on extant taxa to ensure that the selected traits captured significant locomotor variation. The multivariate PGLS fit was then used to run the actual pDFA model (‘mvgls.dfa’ function, ‘mvMORPH’) (Discriminant Function, DF1, Fig. 8; summary statistics in Tables S15-S17). The pDFA model was then tested by predicting the known locomotor categories of extant taxa (‘predict’ R function). Due to low sample size (all analyses being phylogenetically informed, the observations are not represented by single specimens, but by taxa) that could over-estimate the performance of the pDFA, we employed a leave one-out cross-validation (LOOCV). LOOCV involves the exclusion of one random observation from the training set, then employing the pDFA model to predict the class of the excluded observation. This procedure was iterated until all the taxa were excluded once, but avoiding the repeated exclusion of the same taxon, and allowed to compute the misclassification rate (MCR) of the pDFA model. Finally, the pDFA model was applied to subfossil lemurs. To account for uncertainties in the classification of extinct taxa, we followed the approach of Monclús-Gonzalo *et al*. (2023), discouraging the use of posterior probabilities as a reliable proxy for classification uncertainties (e.g. due to overfitting, Qiao *et al*. 2009), and instead using subsampling techniques. Hence, for the subfossil taxa, the locomotor assignment was iterated several times, each time subsampling the training set used to train the pDFA model, and counting the times the extinct species were assigned to each class. As resampling technique, we used LOOCV (detailed above).

**Table 2.** Results from PGLSs and pANOVAs testing the discrimination of ‘SB’ from ‘others’, run with the PC_TP+BM_s (listed in the table, besides the explained portion of variance and by which traits they are mainly driven) that were previously identified as potentially yielding this locomotor information (see Methods for details)

| **Anatomical region**<br>**(articulation)** | **Principal Components** | **Proportion of variance** | **Mainly related to** | **‘SB’-**<br>**‘others’*p*** |
| --- | --- | --- | --- | --- |
| **Proximal femur**<br>**(femoral head)** | PC<sub>TP+BM2</sub> | 19.18% | BV.TV (inversely)<br>DA (inversely) | 0.028 |
| **Distal femur**<br>**(lateral condyle)** | PC<sub>TP+BM3</sub> | 14.34% | DA (inversely) | 0.001 |
| **Distal femur**<br>**(medial condyle)** | PC<sub>TP+BM3</sub> | 14.48% | DA (inversely) | <0.001 |

## Results

The PC1-PC2 scatterplot and variable loading plots deriving from the PCA ran on quantitative locomotor data, revealed four locomotor groups (Fig. 1A): leaping (L), quadrupedal/scrambling/clambering (QSC), suspensory/bridging (SB) and arboreal generalists (AG). Accordingly, we framed in these locomotor categories the taxa for which quantitative data were not available, thus obtaining the final ecological categorization (Fig. 1B) (see further details in Appendix S11). The four categories were not significantly related to BMp (*p*-value_hum_=0.162, *p*-value_fem_=0.476), implying that, in mvPGLS, the BMp–locomotion interaction effects should not be tested and that we could detect distinct effects of body mass and locomotion on trabecular data (Table 1 and below).

### Humeral trabecular structure

The humeral trabecular structure of the extant species did not correlate with locomotion. Indeed, from multivariate PGLSs and phylogenetic MANCOVAs, the humeral datasets were significantly correlated with BMp but not locomotion (Table 1). Also, proximal and distal humerus PCA_TP+BM_ scatterplots and loading vectors showed that PC1_TP+BM_s are primarily related to BMp and to a set of variables—i.e. Tb.Th, Conn.D., Av.Br.Len and BS.TV. The strong body size effect of proximal and distal humeral PC1_TP+BM_s was clear due to the position of great apes and subfossil lemurs (i.e. the largest taxa) on one extreme of PC1_TP+BM_ values with galagids and lorisids (i.e. the smallest taxa) instead lying on the other extreme (Figs. 3-4). In both the humeral epiphyses, correlation tests confirmed the strong BMp effects on PC1_TP+BM_s, showed that Tb.Th, Conn.D., Av.Br.Len and BS.TV significantly contribute to PC1_TP+BM_s (Table S18) and that these trabecular traits are related to BMp themselves (Table S19). Both proximal and distal humeral DA and BV.TV, instead, arose as not significantly related to the respective PC1_TP+BM_s (with the only exception of distal humeral PC1_TP+BM_-BV.TV, but with R^2^=0.29, Table S18) and to body mass (Table S19) (Figs 3-4).

**Fig 4.**
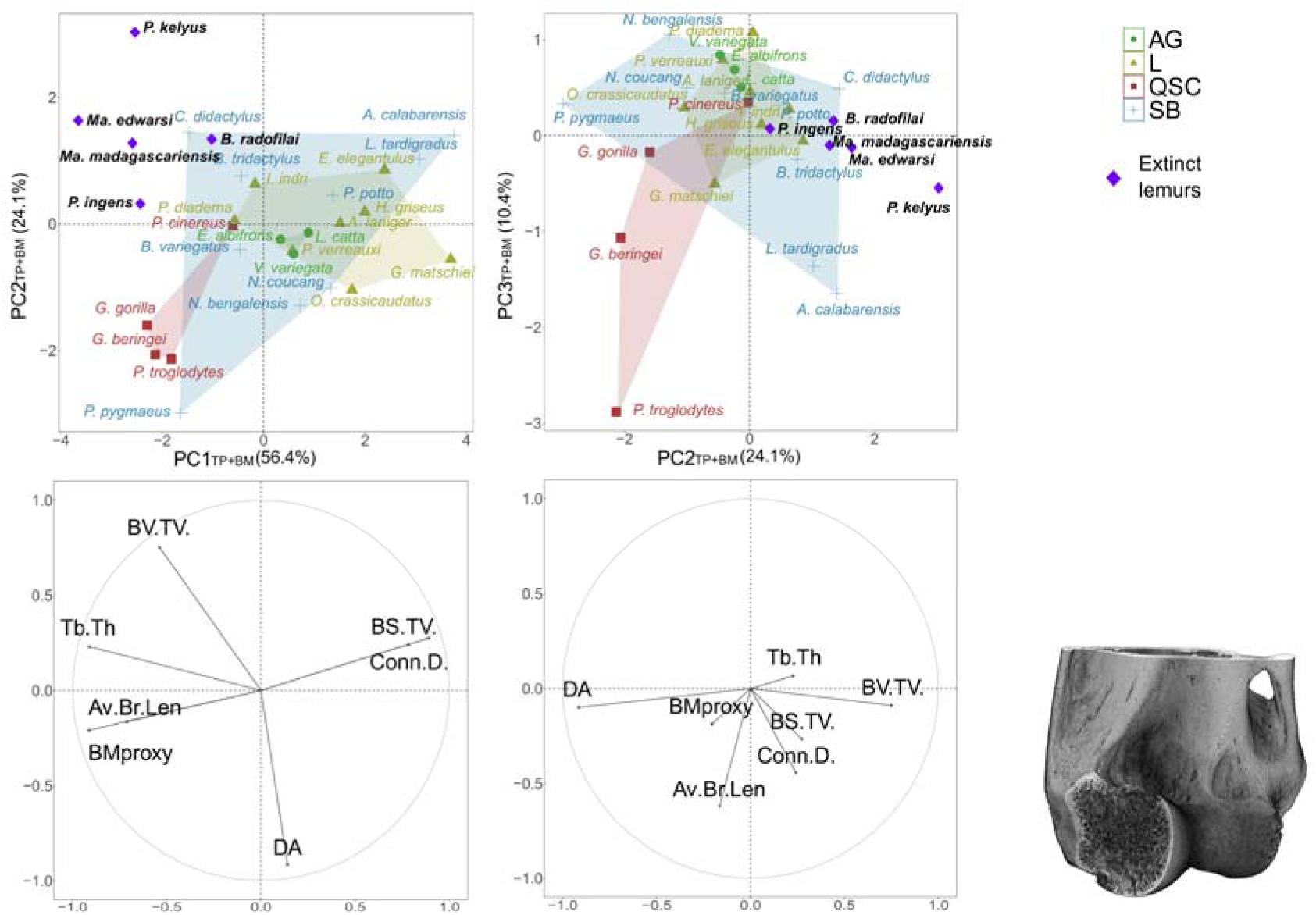
Distal humerus PCA_TP+BM_ results. Top: scatterplots with extant species grouped by locomotor category and subfossil lemur data shown with purple rhombi. Bottom: corresponding variable loading plots.

Humeral proximal and distal PC2_TP+BM_s and PC3_TP+BM_s, instead, do not show any significant relationship with BMp (Figs. 3-4, Table S18). Loading vectors suggest that the PC2_TP+BM_s of both the proximal and distal humerus include information mainly related to DA and BV.TV (that do not contribute to PC1, see above) (Figs. 3-4). It is confirmed by DA-PC2_TP+BM_s and BV.TV-PC2_TP+BM_s correlation tests, combined with the absence of significant correlation tests between any other trabecular variable and PC2_TP+BM_s. Proximal and distal humeral PC3_TP+BM_s contain information related to Av.Br.Len (precisely to its portion of variance not related to BMp, differently from the one driving PC1, see above), since this parameter, yielded the longest vector oriented toward PC3_TP+BM_s (Figs. 3-4) and showed significant correlation with PC3_TP+BM_s (Table S18). Although a few other variables occasionally showed significant correlation with PC3_TP+BM_s too (DA in the proximal, and Conn.D in the distal humerus), their quite low R^2^ (0.13 for DA, 0.20 for Conn.D, Table S18) led us to consider their contribution on the respective PC3_TP+BM_ as minor. The humeral PC2_TP+BM_-PC3_TP+BM_ scatterplots yield a strong overlap of locomotor categories (especially clear for distal humeral data, with almost all the taxa clustering in the same region of the morphospace, i.e. positive PC3_TP+BM_ scores and PC2_TP+BM_ scores ranging between 0 and 1; Figs. 3-4). This, in addition to the non-significant relationship between PC1-PC4 and locomotion from multivariate PGLSs and phylogenetic MANCOVAs (Table 1), led us to not use humeral trabecular properties in the following steps of the analysis.

In the humeral PC2_TP+BM_-PC3_TP+BM_ morphospaces (in which allometric effects are minimized, see above), subfossil lemurs overall cluster in the same regions (i.e. positive PC2_TP+BM_-negative PC3_TP+BM_ scores for the proximal humerus, positive PC2_TP+BM_ scores for the distal humerus). In the sloth-lemurs’ range of variation, *Palaeopropithecus* spp. tend to occupy the most peripheral position (with the exception of *P. ingens* in the distal humerus), *Babakotia* lies the most closely to the other extant taxa, and *Mesopropithecus* (data available only for the proximal humerus) lies in another peripheral region of the morphospace (close to *A. calabarensis*) (Figs. 3-4). *Megaladapis* spp. tend to lie close to *Babakotia*. Interestingly, in the humeral PC2_TP+BM_–PC3_TP+BM_ morphospaces, most subfossil lemurs tend to occupy distinct regions, i.e. they do not overlap with those of extant mammals. Indeed, in the proximal humerus, *Palaeopropithecus* spp., *Me. dolichobrachion*, and *Me. madagascariensis* lie outside the convex hulls of extant mammal locomotor categories (Fig. 3). In the distal humerus, a similar pattern is observed for *P. kelyus* and *Ma. edwardsi*, while *Babakotia* and *Ma. madagascariensis* occupy peripheral positions within the SB area and trend toward *P. kelyus* and *Ma. edwardsi* (Fig. 4). Both the proximal and distal humeral outlying pattern of subfossil lemurs are mainly driven by high BV/TV (Figs. 3-4).

### Femoral trabecular structure

Femoral data yielded patterns related to locomotor categories. Indeed, trabecular data from lateral and medial condyle proved to be significantly related to locomotion, as shown by multivariate PGLSs and phylogenetic MANCOVAs (Table 1). These epiphyseal regions, such as the proximal femoral, are significantly related to body mass too (Table 1) but, since locomotor and allometric effects are separated in PGLSs (see above), lateral and medial condyle size-corrected data are significantly related to locomotor categories. Although not reaching significance, proximal femoral data yielded a particularly low *p* for a relationship with locomotion (0.09, Table 1).

PC1_TP+BM_s across three femoral datasets are explained by body mass and collects the allometric effects, as shown by the position of great apes/subfossil lemurs (i.e. the largest taxa) and galagids/lorisids (i.e. the smallest taxa) at the two extremes of PC1_TP+BM_ scores. It is also suggested by BMp loading vectors’ directions (Figs. 5-7), additionally indicating that PC1_TP+BM_s are driven by a portion of variance of a subset of trabecular variables that are likely including size-effects themselves. Through correlation tests across the femoral datasets, we confirmed the significant relation between PC1_TP+BM_s and BMp, we identified the trabecular variables driving PC1_TP+BM_s, i.e. Tb.Th., Conn.D., BS.TV and Av.Br.Len (Table S18), and we found that they drive PC1_TP+BM_ through their relationships with BMp (Table S19). In the proximal femur and in the medial condyle, a portion of BV.TV variance too is correlated with PC1_TP+BM_ but with quite low strength (R^2^_prox_=0.24, R^2^_med.con_=0.24, Table S18). Likewise, BV.TV significantly but quite weakly relates to BMp in these two levels (R^2^_prox_=0.21; R^2^_med.con_=0.23, Table S19). DA minorly contributes to femoral PC1_TP+BM_s, since it does not correlate or weakly correlates with PC1_TP+BM_s (R^2^_lat.con_=0.14) (Table S18) and does not correlate with BMp (Table S19).

**Fig 5.**
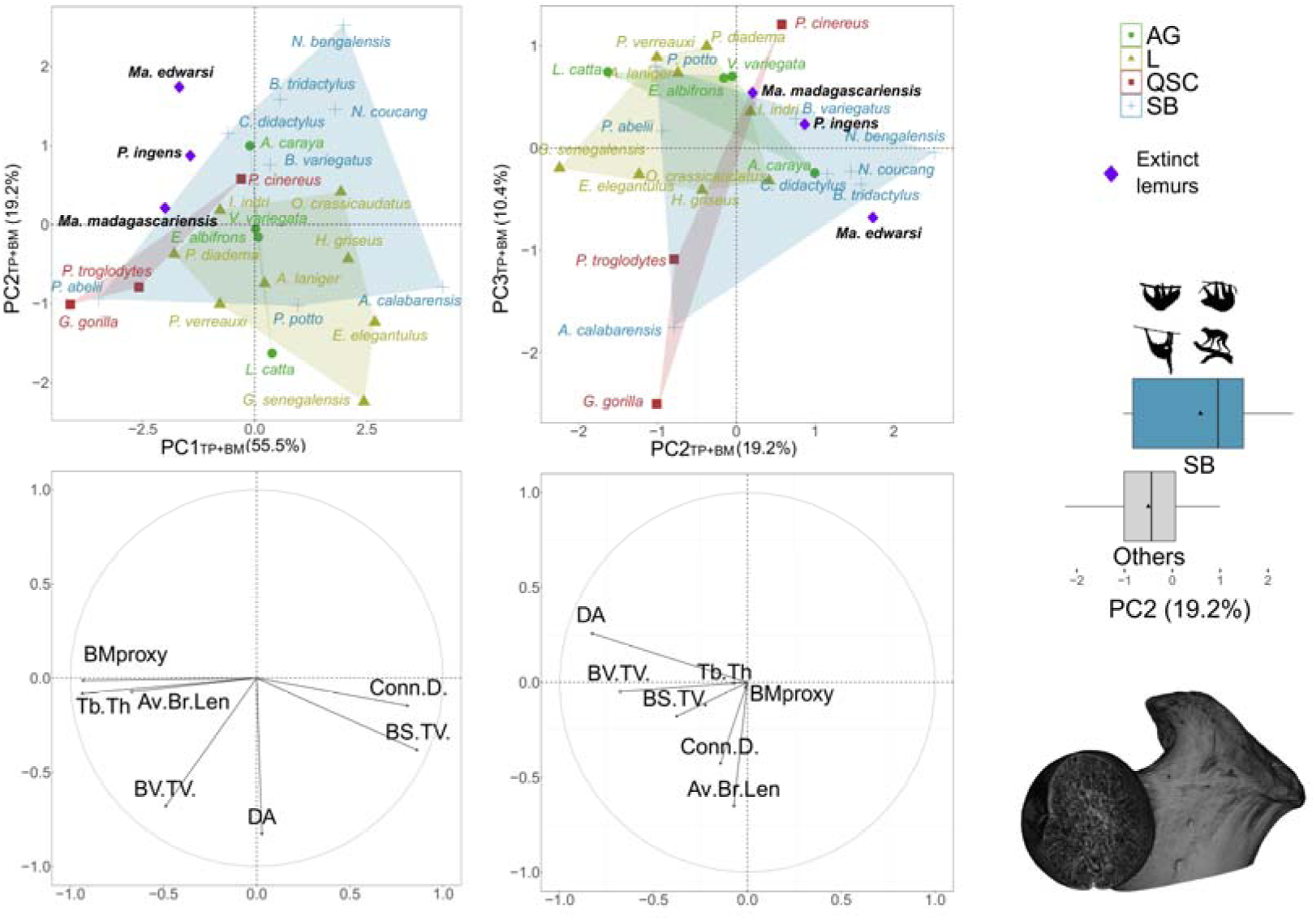
Proximal femur PCA_TP+BM_ results. Top: scatterplots with extant species grouped by locomotor category and subfossil lemur data shown with purple rhombi. Bottom: corresponding variable loading plots. Middle right: univariate distribution of the PC_TP+BM_ along which we identified the separation of ‘SB’ from ‘others’.

Femoral PC2_TP+BM_s and PC3_TP+BM_s do not include size effects, as suggested by BMp vectors in the loading plots (Figs. 5-7) and PC2_TP+BM_s and PC3_TP+BM_s not correlating with BMp (Table S18). Correlation tests show that proximal femoral PC_TP+BM_2 is mainly driven by DA, especially (R^2^=0.69), and BV.TV (R^2^=0.47) and no other variables (Table S18). Both DA and BV.TV contribute negatively to proximal femoral PC2_TP+BM_, i.e. higher PC2_TP+BM_ scores are yielded for lower DA and BV.TV (Fig. 5, Table 2). Correlation tests also tell us that lateral and medial condyle PC2_TP+BM_s are driven by BV.TV primarily, less strongly by Av.Br.Len and not by other variables (no *p*<0.05, with the only exception of DA in the medial condyle, *p*=0.02, but with weak correlation, R^2^=0.19) (Table S18). In the proximal femur, PC3_TP+BM_ captures information on Av.Br.Len and, but quite weakly, on Conn.D (R^2^= 0.18), and no other variables (Fig. 5, Table S18). Correlation tests show that lateral and the medial condyle PC3_TP+BM_s are mainly driven by DA and no other variables (Table S18) (Figs. 6-7). DA contributes negatively to lateral and medial condyles PC3_TP+BM_s, i.e. lower DA is shown by taxa yielding higher PC3_TP+BM_ scores (Figs. 6-7, Table 2).

**Fig 6.**
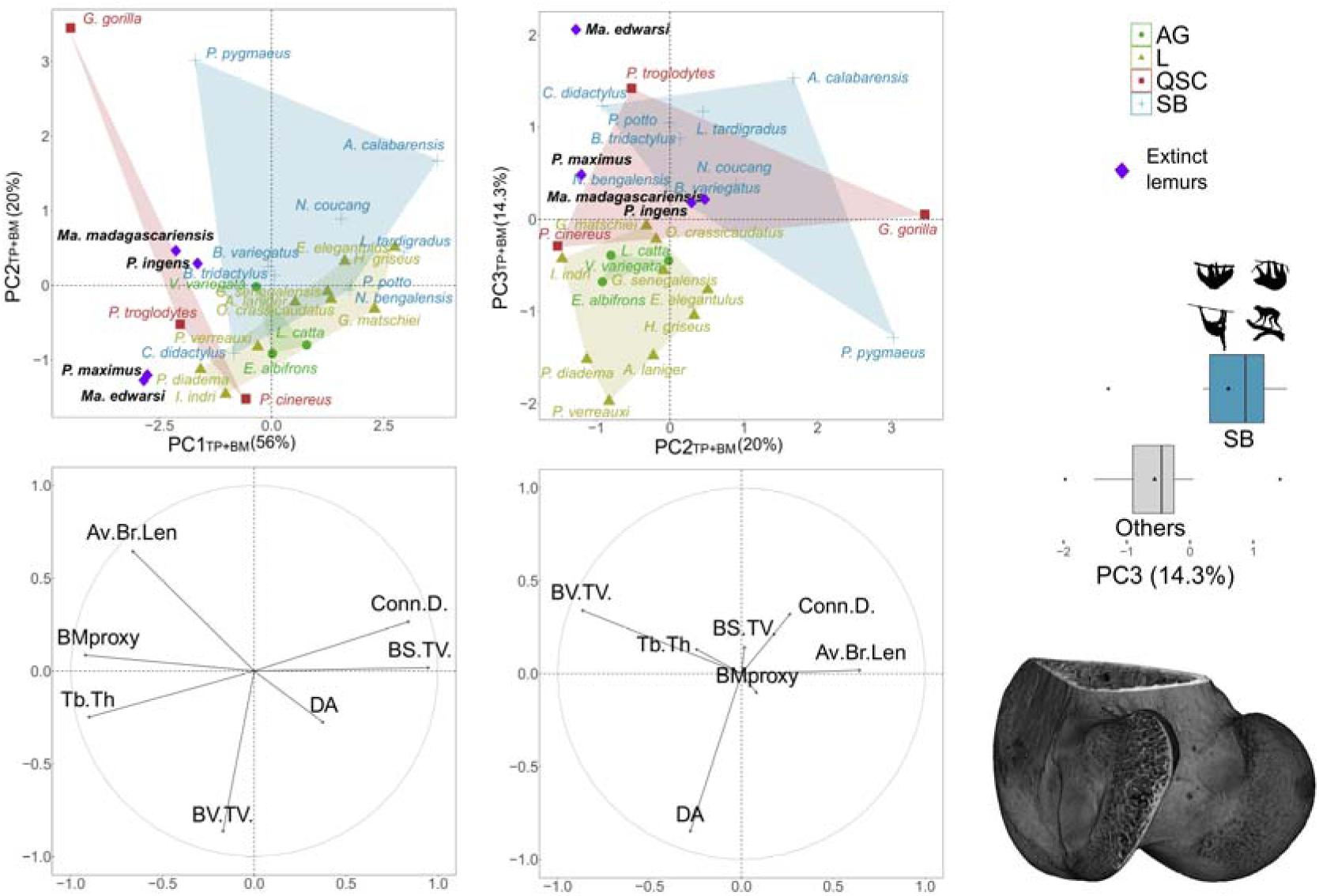
Lateral femoral condyle PCA_TP+BM_ results. Top: scatterplots with extant species grouped by locomotor category and subfossil lemur data shown with purple rhombi. Bottom: corresponding variable loading plots. Middle right: univariate distribution of the PC_TP+BM_ along which we identified the separation of ‘SB’ from ‘others’.

**Fig 7.**
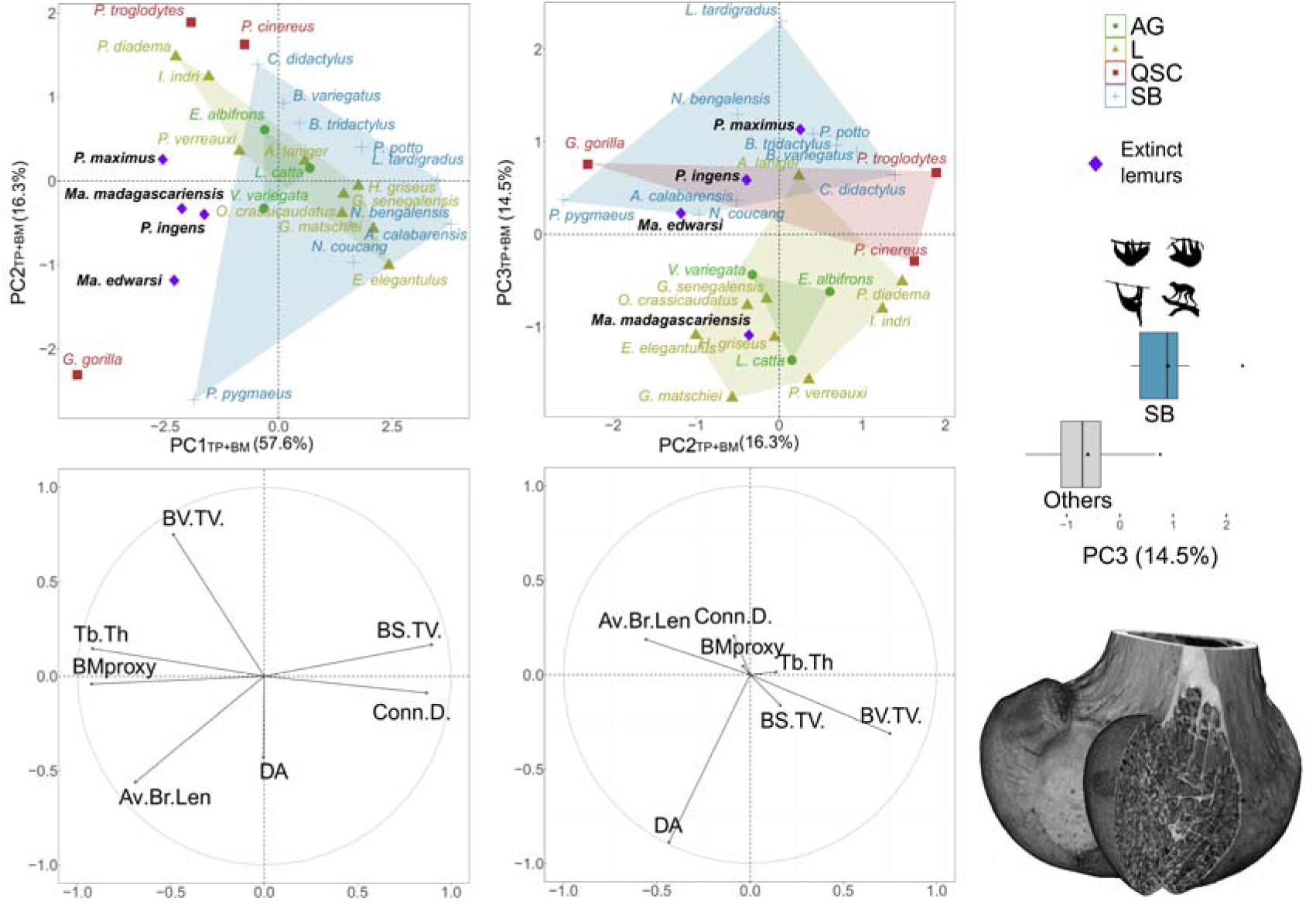
Medial femoral condyle PCA_TP+BM_ results. Top: scatterplots with extant species grouped by locomotor category and subfossil lemur data shown with purple rhombi. Bottom: corresponding variable loading plots. Middle right: univariate distribution of the PC_TP+BM_ along which we identified the separation of ‘SB’ from ‘others’.

**Fig 8.**
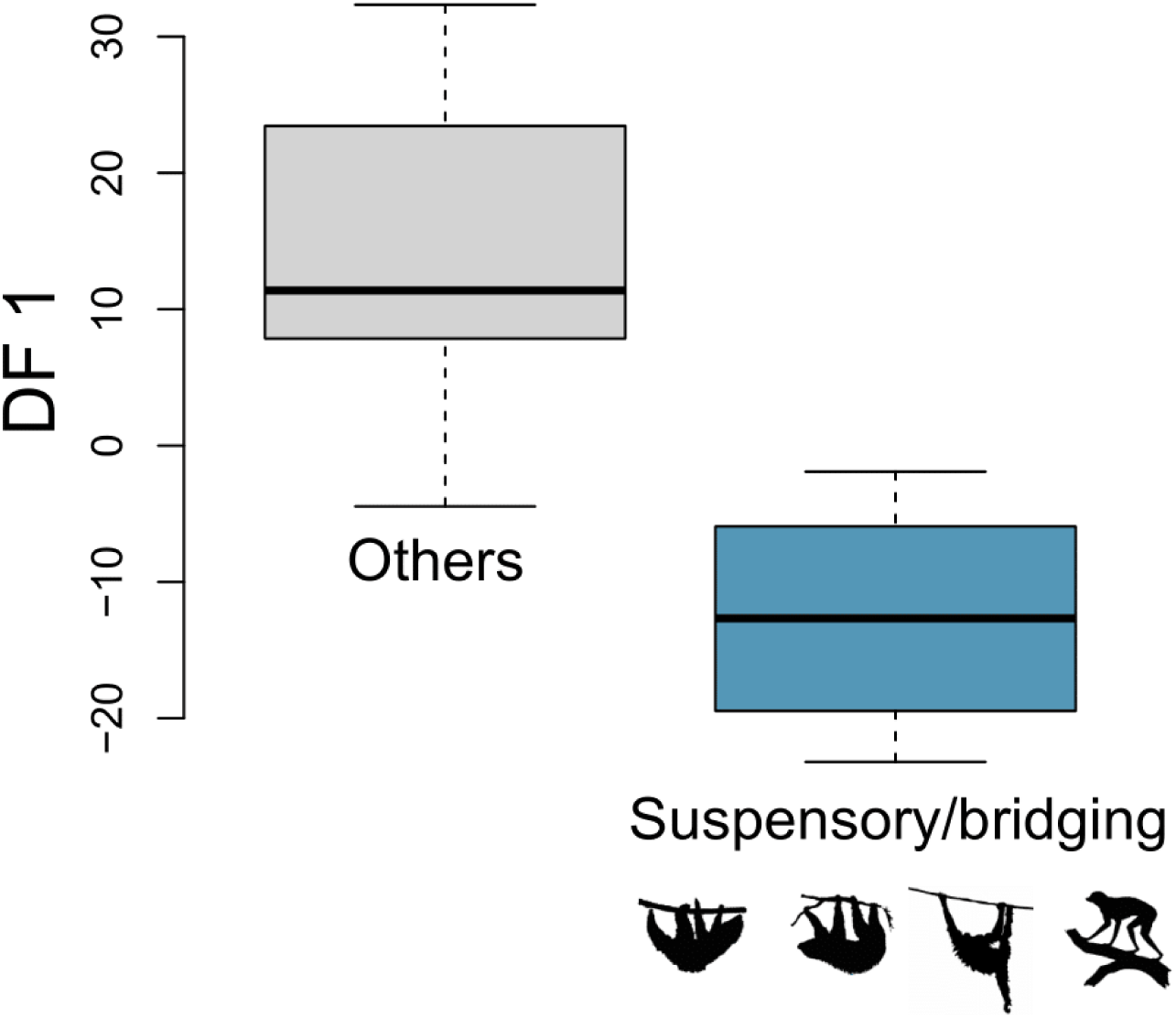
Distribution of DF1, deriving from a pDFA ran on four predictor trabecular variables.

On the femoral PC2_TP+BM_-PC3_TP+BM_ morphospaces, we visually identified a pattern of discrimination of ‘SB’ from all the other taxa (‘others’ hereafter) that possibly caused the significant relationship with locomotion arising from multivariate PGLSs and phylogenetic MANCOVAs for two femoral epiphyseal regions (Table 1, see also above). We focused on the discrimination between SB and ‘others’, because in all the femoral anatomical levels, subfossil lemurs tended to lie in the morphospace areas occupied by ‘SB’, making this trend of interest for the focus of this work (Fig. 5-7). In the proximal femur, ‘SB’ tend to yield high PC2_TP+BM_ scores (although *Pe. potto*, *Po. abelii* and *A. calabarensis* lie within ‘others’, Fig. 5). In the lateral and medial condyle, ‘SB’ yielded higher PC3_TP+BM_ scores (with some exceptions: for the lateral condyle, the ‘SB’ *Po. pygmaeus* lies within ‘others’, and *Pa. troglodytes*, belonging to ‘others’, instead lies within ‘SB’; for the medial condyle, the position of few taxa belonging to ‘others’, i.e. *G. gorilla, A. laniger, P. troglodytes*, cause overlap with the ‘SB’ range). The PC2_TP+BM_ pattern for the proximal femur and the PC3_TP+BM_ patterns for lateral and medial condyles were all confirmed qualitatively through PGLSs and ANOVAs (Table 2).

### Phylogenetic discriminant function analysis

As detailed above, the discrimination between ‘SB’ and ‘others’ was the only pattern of interest yielded by trabecular structure. It was only caused by femoral traits and, specifically, by higher PC2_TP+BM_ (primarily corresponding to lower BV.TV and DA) in the femoral head PCA_TP+BM_ (Fig. 5) and by higher PC3_TP+BM_ (mainly corresponding to lower DA) in both the lateral and the medial condyle PCA_TP+BM_s (Fig. 6-7). Hence, we took the trabecular traits primarily contributing to these femoral PC_TP+BM_s (i.e. BV.TV_prox_, DA_prox_, DA_lat.con_, DA_med.con_, identified as detailed above) and we built a multivariate heterogeneous dataset only including extant taxa (i.e. the training set) on which we trained a pDFA model that is, accordingly, potentially able to classify taxa in ‘SB’ or ‘others’. The heterogeneous dataset confirmed the potential to discriminate ‘SB’ from ‘others’ (multivariate PGLS and phylogenetic MANOVA, *p*= 0.002) and the pDFA model trained on the training set yielded a MCR of 0.14. We considered this value as sufficiently low (i.e. the model correctly predicted locomotion for the 86% of the observations excluded during LOOCV). The pDFA generated a single Discriminant Function, i.e. DF1, since the model accounted for only two classes, i.e. ‘SB’ vs. ‘others’. DF1 was positively correlated with all the variables on which the model was run (Table S16) and, as shown with a boxplot, it neatly separated ‘SB’ (lower DF1, i.e. thus lower femoral BV.TV_prox_, DA_prox_, DA_lat.con_ and DA_med.con_) from ‘others’ (Fig. 8). The pDFA model, run on *Megaladapis madagascariensis*, *Megaladapis edwarsi* and *Palaeopropithecus ingens* (the three subfossil lemur species for which we had data coming from proximal femur, lateral and medial condyle), classified *M. edwarsi* and *P. ingens* as ‘SB’ in the 100% of the iterations, while *M. madagascariensis* was never classified as ‘SB’.

## Discussion

### Locomotor signal in extant mammal trabecular structure

When assessing the locomotor categories that we established mapped onto the phylogeny it becomes evident that, although some clades are characterized by displaying a single locomotor category (e.g. all galagids are classified into the leaping category), the overall distribution reveals multiple independent acquisitions of the same (broadly defined) locomotor behavior across distant lineages (e.g., SB in *Pongo*, Bradypodidae, Choloepodidae, and Lorisidae; leaping in Galagidae, Indriidae, and *H. griseus*) (Fig. 1B). This distribution supports the interpretation that any trabecular similarity among taxa sharing the same locomotor category in different regions of the phylogeny may be attributed to functional adaptations related to locomotor behaviors. Also, the relationship between trabecular structure and locomotion was tested through phylogenetically informed analyses (e.g. PGLS, pDFA), that account for shared phylogenetic history of the studied taxa. It ensures that the locomotion-related patterns that we found and discuss below are not due to phylogenetic affinity. We aimed not only to exclude phylogenetically driven patterns but potentially to observe them, in case they were dominant (i.e. by observing closely related taxa clustering in a region of the morphospace, e.g. subfossil lemurs more closely resembling other Madagascar strepsirrhines). This rationale was at the base of our choice to use traditional PCAs instead of other techniques that erase phylogenetic trends (e.g., phylogenetic PCA; Polly *et al*. 2013), for qualitative assessments.

As for allometric effects, in both humeral and femoral PCAs, they are mostly captured in PC1_TP+BM_s (Figs. 3-7). Since PC1 normally accumulates most of the variance in a PCA, a strong allometric effect on PC1_TP+BM_s suggests that body mass is a major driving factor of trabecular bone in the sample studied. This was also evident from PGLSs (Table 1) and it is consistent with previous findings in mammals (Amson *et al*. 2017; Doube *et al*. 2011), especially primates (Alfieri *et al*. 2025a; Ryan and Shaw, 2013), but also other amniotes—both broad amniote samples (e.g., Gônet *et al*. 2023) or in specific clades as birds (Alfieri *et al*. 2025b; Doube *et al*. 2011) and non-avian reptiles (Plasse *et al*. 2019). Investigating detailed scaling patterns is out of the scope of this work, but it is interesting to note that, for instance, Conn.D and BS/TV positively co-vary in driving PC1_TP+BM_s across all the studied regions (see loading vectors’ directions in Figs. 3-7), which is consistent with previous works (e.g. Gônet *et al*. 2023) and expected structural patterns (i.e. the higher Conn.D., the higher the trabecular number, the greater the bone surface).

In all the PGLSs that we run across the studied anatomical regions, effects of body mass were separated from locomotor ones. Hence, the effects of locomotion found in the PGLSs (Table 1) were already isolated from allometry. Also, all the locomotor patterns discussed below were derived from PC2_TP+BM_s or PC3_TP+BM_s and from two variables, i.e. BV/TV and DA, that largely did not include body mass effects. Indeed, no significant correlation with BMp was found for these parameters in the femoral datasets, with the only exceptions of BV/TV in the head and lateral condyle that weakly correlates to BMp (Table S19), and with BV/TV from lateral condyle that, importantly, was not used in the locomotor reconstructions. DA and/or BV/TV not being correlated to body mass aligns with many previous works (e.g. Barak *et al*. 2013; Cotter *et al*. 2009; Doube *et al*. 2011; Fajardo *et al*. 2013; Gônet *et al*. 2023; Kivell *et al*. 2018; Plasse *et al*. 2019; Rolvien *et al*. 2017) and indicates that the locomotion-related patterns we propose are independent of allometric effects. Moreover, the successful isolation of these two allometrically-independent variables supports our choice to account for body mass effects by excluding PC1s (common practice in shape analyses, e.g. Klingenberg, 2022; Somers, 1989) instead of using other size-correction techniques, such as those based on regression residuals (which previously proved to not fully exclude allometric effects from trabecular variables; e.g. Alfieri *et al*. 2025a).

The femoral trabecular structure yielded a relationship with locomotor behavior, i.e. significant correlation between femoral condyles’ trabecular parameters and locomotor categories (Table 1, with proximal femoral trabecular properties yielding a *p* approaching significance). PC2_TP+BM_-PC3_TP+BM_ morphospaces suggested that femoral trabecular structure is overall able to discriminate suspensory/bridging mammals (SB, i.e. lorisids, tree-sloths and orangutans) from all the other extant mammals. Specifically, SB are distinct for showing averagely higher proximal femur PC2_TP+BM_ scores (Fig. 5), higher lateral condyle PC3_TP+BM_ scores (Fig. 6) and higher medial condyle PC3_TP+BM_ scores (Fig. 7). These patterns were confirmed quantitatively (see also boxplots in Figs. 5-7) and—as indicated by loading vectors—are primarily driven by low BV/TV_prox_, DA_prox_, DA_lat.con_ and DA_med.con_ (Figs. 5-7, Table 2). Accordingly, when these four femoral trabecular parameters were analysed, they significantly discriminated SB from other taxa and, when used to run a pDFA, yielded a DF that neatly discriminates SB from other mammals (Fig. 8) and that correctly classifies them with a high success rate (i.e. 86%). Thus, multiple lines of evidence suggested that SB differ from other mammals by exhibiting lower bone volume (i.e. low BV/TV) in the proximal femur and a more isotropic trabecular structure (i.e. low DA) across all femoral levels. Proximal femur trabecular structure proved to be related to locomotor activities in many previous studies mostly focusing on mammal— especially primate—femoral head (e.g. Alfieri *et al*. 2022; Fajardo and Müller, 2001; Georgiou *et al*. 2020, 2019; MacLatchy and Müller, 2002; Mielke *et al*. 2018; Ryan and Ketcham, 2005, 2002b; Ryan and Krovitz, 2006; Ryan and Shaw, 2012; Saparin *et al*. 2011; Tsegai *et al*. 2018), aligning with our results.

#### Femoral low DA and BV/TV in suspensory/bridging mammals: functional interpretations

A functional interpretation for lower femoral DA and BV/TV in SB compared to other mammals is consistent with the fact that these two parameters can generally provide sufficient information on the biomechanical regime, and accordingly, on locomotor habits, since they cumulatively explain up to 98% of bone stiffness (through their contribution to Young’s modulus; Maquer *et al*. 2015; Stauber *et al*. 2006).

As shown experimentally, DA tends to follow loading directions (Biewener *et al*. 1996; Pontzer *et al*. 2006), which may be in turn informative on locomotor habits. DA captures the degree of preferential alignment of trabeculae, hence not the specific directions (e.g. in degrees of arc) of trabeculae but the extent to which they are uniformly oriented (as computed here, from DA=0, i.e. highly multidirectional or fully isotropic structure, to DA=1, highly directional or fully anisotropic structure; Harrigan and Mann, 1984). Thus, DA positively relates to directional stereotypy of movements involved in locomotor activities—e.g. high DA is caused by fossoriality (Amson *et al*. 2017) and bipedalism (Georgiou *et al*. 2019; Ryan *et al*. 2018)—while low DA derives from multidirectional locomotor repertoires. Arboreality represents a paradigmatic example of the latter condition, since moving in trees results in highly variable directions of biomechanical loadings (Carlson, 2005; Demes *et al*. 2006). Our sample is represented by taxa that, to several extents, show arboreal habits (Fig. 1), hence lower DA in both proximal and distal femur of SB possibly reflect their outstandingly broader mobility in hip and knee joints, even compared to other arboreal taxa which joints are expected to be relatively mobile too. An exception is represented by leaping taxa that, although being fully arboreal, are characterized by a distinctive behavior related to powerful leaps and take-offs that cause high directional stereotypy and, as expected, high DA (as found by Ryan and Ketcham, 2005, 2002b). An extreme mobility in hip and knee joints—related to suspensory and/or bridging locomotion—has been reported, or derived from joint/muscle configuration, for orangutans (Thorpe and Crompton, 2005; Zihlman *et al*. 2011), lorisids (Ishida *et al*. 1992; Runestad, 1997; Youlatos *et al*. 2025) and tree-sloths (Marshall *et al*. 2021; Nyakatura, 2012). Thus, SB taxa possibly occupy a positive extreme on the arboreal range of hindlimb joint mobility, and the consequent extraordinarily multidirectional loadings acting on the hip and knee result in particularly isotropic trabeculae, i.e., low DA.

BV/TV measures the proportion of inner epiphysis occupied by ossified fraction and informs on trabecular bone compactness. Increased loadings are expected to unbalance bone modeling (*sensu* Barak, 2019) in favor of bone deposition at the expense of bone reabsorption and resulting in a greater bone fraction, while the opposite is expected under decreased loading regimes (Pearson and Lieberman, 2004). Within an epiphysis, it is reflected by BV/TV and, accordingly, a positive relationship between trabecular fraction and loadings has been found or hypothesized (e.g. Chang *et al*. 2008; Modlesky *et al*. 2008; Ryan and Shaw, 2015). In this framework, the lower femoral BV/TV shown by SB is possibly justified by particularly low hindlimb loadings related to arboreality based on suspension and bridging. This condition has been consistently discussed for tree sloths—due to their outstandingly cautious lifestyle (e.g. Alfieri *et al*. 2022, 2021)—lorisid—hence, considered remarkable exceptions to the primate hindlimb dominance (see above) (Hanna *et al*. 2017; Ishida *et al*. 1990; Yapuncich and Granatosky, 2021)—and orangutans, accordingly showing poorly robust hindlimb diaphyses (e.g. Ruff, 1988). Based on these functional interpretations, we posit that low DA and BV/TV in the respective femoral epiphyses of subfossil lemurs, is a clear indicator of suspensory/bridging in these extinct taxa.

### The locomotor behavior of palaeopropithecids

We found strong confirmation for *Palaeopropithecus* as the most extremely suspensory sloth-lemur. Indeed, *P. ingens* and/or *P. maximus* always felt within the range that we interpreted as characterizing SB (i.e. higher femoral head PC2 scores and femoral condyles PC3 scores, Figs. 5-7). Furthermore, *P. ingens*, the only sloth-lemur species for which we had data coming from all the studied femoral epiphyseal regions, was classified as SB in 100% of the iterations by the pDFA model. Thus, we add femoral trabecular structure to the growing list of anatomical traits (e.g. Godfrey *et al*. 1995; Godfrey and Jungers, 2003; Granatosky, 2018, 2020, 2022; Hamrick *et al*. 2000; Jungers, 1980; Jungers *et al*. 1991; Marchi *et al*. 2016; Patel *et al*. 2013; Shapiro *et al*. 1994, 2005; Wunderlich *et al*. 1996) that now with very high confidence indicate that *Palaeopropithecus* spp. were large lemurs exhibiting highly specialized suspensory arboreality.

Previous works have proposed suspensory habits in *Babakotia* and *Mesopropithecus* too, but less extremely compared to *Palaeopropithecus* (Marchi *et al*. 2016 and references therein). Our results do not allow us to contribute to this view, due to the availability/preservation of the specimens and the informative power that arose from the study of extant taxa, i.e. only femoral trabecular structure can be used for locomotor inference. Indeed, the fact that we only had humeral data for *Babakotia* and *Mesopropithecus* limits our inferential power on these smaller sloth-lemurs. Thus, we encourage further works to study femoral trabecular data on *Babakotia* and *Mesopropithecus* to test if their DA and BV/TV are low enough to indicate suspensory/bridging habits for these two taxa too.

### The locomotor behavior of *Megaladapis* spp

Despite early reconstructions not necessarily inferring an arboreal lifestyle for *Megaladapis*— with even an aquatic ecology having been suggested following the first fossil discoveries (Godfrey and Jungers, 2002 and references therein)—in the last decades, evidence from many postcranial skeletal traits indicate an arboreal lifestyle for koala-lemurs (Godfrey *et al*. 2016; Godfrey and Jungers, 2002; Jungers, 1978; Wunderlich *et al*. 1996). However, doubts remain concerning the type of arboreal behavior koala-lemur engaged in across species. According to an earlier view, all the species belonging to the *Megaladapis* genus engaged in vertical climbing and clinging, Hence, in this framework, *Megaladapis* spp. would be represented by size and geographic variants (Carleton, 1936; Jungers, 1977, 1978). However, more recent inferences hypothesized the presence of two sub-lineages: *Megaladapis*—including the medium-small sized species *M. madagascariensis* and *M. grandidieri*, widely distributed in Madagascar, exhibiting more arboreal habits and mainly featuring vertical climbing (with additional potential behaviours as quadrupedal hanging and pedal suspension, Jungers *et al*. 2008, 2002b; Wunderlich *et al*. 1996, 1994)—and *Peloriadapis*—corresponding to *M. edwarsi* that, mainly due to its large body size, has been proposed as relatively more terrestrial compared to other *Megaladapis* species (Jungers *et al*. 2008, 2002b; Wunderlich *et al*. 1996, 1994). Our results did not yield the informative power allowing us to identify vertical climbing from femoral trabecular bone. This locomotor habit can be ascribed to the arboreal generalist (AG) and/or quadrupedalism/scrambling/clambering (QSC) within our categories (especially the latter, remarkably representing the koala *Phascolarctos cinereus*), but the trabecular data that we analyzed did not reveal differences related to these locomotor styles. Instead, our results provide support for the existence of locomotor differences between small/medium and large koala lemurs. Indeed, focusing on the femoral data, the smallest (*M. madagascariensis*) and the largest (*M. edwarsi*) koala lemurs are quite distant in the femoral morphospaces, with *M. edwarsi* lying within the SB range of variation (i.e. higher PC2_TP+BM_ scores in the head, Fig. 5; higher PC3_TP+BM_ scores in the medial condyle, although close to the ‘non-SB’ range, Fig. 7) or even yielding the most extreme score in the direction interpreted as characterizing SB (i.e. extremely high PC3_TP+BM_ score in the lateral condyle, Fig. 6). In this regard, the result of pDFA for *M. madagascariensis* and *M. edwarsi* are striking. The smaller taxon *M. madagascariensis* was never classified as ‘SB’ across all the iterations, while the larger *M. edwarsi* was classified as SB in 100% of the iterations, mirroring *Palaeopropithecus* (see above). The remarkable aspect of this outcome mainly concerns the reconstruction of *M. edwarsi* as an extremely suspensory/bridging *Palaeopropithecus*-like taxon. Indeed, it represents a novel view that, to the best of our knowledge, has never been proposed before. While in the traditional locomotor reconstruction of koala-lemurs, quadrupedal hanging and pedal suspension have been proposed in addition to the dominant vertical climbing (Jungers *et al*. 2008, 2002b; Wunderlich *et al*. 1996, 1994), this description did not refer to interspecific differences within *Megaladapis*, i.e. the wide arboreal repertoire was proposed for the entire genus. Here, we instead highlight intrageneric locomotor differences in *Megaladapis* but not aligning to the previously proposed relatively higher terrestriality of *M. edwarsi* (Jungers *et al*. 2008, 2002b; Wunderlich *et al*. 1996, 1994). Here, the latter instead appears to be highly adapted to suspensory/bridging arboreal habits—an adaptation that involves greater exploitation of trees, rather than reduced use. We are aware that this is a preliminary result since it should be confirmed by future work representing more *M. edwarsi* data and more directly testing this hypothesis (e.g. studying koala-lemurs femoral diaphyseal cross-sections) and due to the contradictory humeral results. However, *M. edwarsi* being a fully suspensory/bridging arboreal species is remarkable considering the reconstructed body mass for this taxon, ca. 85 kg (Jungers *et al*. 2008). As of today, orangutans are the largest known extinct or extant mammals showing suspensory/bridging arboreal habits, with adult males of the largest orangutan species—i.e. Bornean orangutan *P. pygmaeus*—reaching 74 ± 9.78 kg (Rayadin and Spehar, 2015)—estimates which are consistent with older body mass estimates (e.g. 78 kg; Smith and Jungers, 1997). Other paradigmatic arboreal suspensory/bridging mammals are characterized by smaller body mass (e.g. tree sloths, 4-6 kg, Nowak, 1999), even when other possibly extremely suspensory/bridging large-sized extinct lemurs are considered (*Palaeopropithecus* spp., ca. 41-46 kg, Jungers *et al*. 2008). Thus, the extreme suspensory/bridging adaptations in *M. edwarsi* would push the boundary or at least achieve the one of Bornean orangutans concerning the largest mammal capable of this behavior, along with all the distinctive morpho-physiological adaptations required to sustain such a peculiar lifestyle (Granatosky, 2022; Nyakatura *et al*. in press, and references therein). *Archaeoindris* is another sloth-lemur (not studied in this work) and it is the largest reconstructed subfossil lemur, approaching the size of a male gorilla, i.e. ca. 161 kg (Jungers *et al*. 2008). However, differently from other sloth-lemurs such as *Palaeopropithecus*, its locomotion is still poorly understood due to the few fragmented preserved postcranial elements for which locomotor inferences have been attempted: e.g., slow-moving arboreality, arboreal climbing and clinging, or ground sloth-like scansoriality (Godfrey *et al*. 2016; Godfrey and Jungers, 2002). Retrieving and studying additional postcranial material of *Archaeoindris* may help identify new instances of exceptionally large mammals with varying degrees of arboreal adaptation.

### Final remarks

The absent locomotor signal in humeral trabecular bone, as revealed by multivariate inferential analyses (Table 1) and the fact that no visual distinctions between locomotor categories could be found in PC2_TP+BM_-PC3_TP+BM_ morphospaces (Figs. 3-4), contrasts the femoral trabecular pattern and, in this way, aligns to previous evidence. A discrepant pattern between humeral and femoral anatomy, with the latter being more informative on locomotor habits, arose in Ryan and Shaw 2012, agreeing with outcomes from primate diaphyseal principal moments of area (Carlson, 2005). Also, Ryan and Walker (2010) highlighted different patterns between primate humeral and femoral trabecular bone. These interlimb trends were previously justified with the fact that in primates the vertical peak ground reaction forces to which hindlimbs are subjected are higher than those experienced by forelimbs (Scherf *et al*. 2013). This evidence is consistent with indications of higher hindlimb loadings provided by Ryan and Walker (2010) and has been built by studying different quadrupedal species from almost all primate sub-groups (e.g. Demes *et al*. 1994; Kimura, 1992, 1985; Kimura *et al*. 1979; Schmitt and Hanna, 2004; Young, 2012). Hence it may be considered a general primate condition (‘hindlimb dominance’) that is opposite to the one shown by other mammals (Larson, 1998; Young, 2012). If we consider that our extant mammal sample is largely composed of primates, hindlimb dominance may potentially explain the discrepant results between femoral and humeral trabecular structure. The composition of our sample may be related to another explanation for the absence of locomotor signal in humeral trabecular bone. Indeed, many studies that found an absent/weak locomotor signal in humeral trabecular structure (see above) focused on distantly related species, representing different phylogenetic sub-groups (primates in Ryan and Shaw, 2012; Ryan and Walker, 2010), in this regard mirroring our sample (Fig. 1). Hence, humeral trabecular structure may not strongly co-vary with locomotor behavior when it is analyzed across phylogenetically broad samples. This condition may not allow us to distinguish even highly divergent locomotor styles (e.g. see the strikingly similar humeral trabecular bone in bipeds and quadrumanous climbers in Ryan and Shaw, 2012). On the other hand, when restricted taxonomic contexts are analyzed, broad scale effects (e.g. phylogenetic, allometric) may be minimized, and functional aspects may be highlighted in forelimb trabecular structure. It would be consistent with the apparent locomotor signal found in humeral trabecular bone when restricted groups, e.g. hominids, were analyzed (Kivell *et al*. 2018; Scherf *et al*. 2013), contrasting results from works focusing on distantly related taxa (Ryan and Shaw, 2012; Ryan and Walker, 2010). A similar explanation has been proposed for the absence of locomotor signal in the trabecular structure of other skeletal regions, again in primates (e.g. distal fibula; Alfieri *et al*. 2025a) and even outside mammals (e.g. squamates, turtles and crocodiles, Plasse *et al*. 2019; extant birds; Alfieri *et al*. 2025b). Future work on humeral vs. femoral trabecular structure of more extensive and diverse samples, may help to elucidate the locomotor effects in taxonomically and phylogenetically broad contexts and how humeral/femoral patterns differ between primates and non-primates.

Apart from these considerations, it should be noted that in the proximal humeral PC2_TP+BP_– PC3_TP+BP_ biplot, *P. pygmaeus* occupies an outlying position compared to the other SB taxa, which tend to show higher PC2 _TP+BP_ scores (Fig. 3). It suggests that the outlying proximal humeral results for this SB species alone may have prevented detecting significant locomotor effects from PGLSs (*p*-value=0.21, Table 1). Remarkably, *P. pygmaeus* proximal humerus data come from a single specimen (i.e. *Pongo pygmaeus* ZSM 1907-609, Table S1), whose patterns might not be representative of the species (a point that future studies may clarify). Thus, proximal humeral trabecular bone might have informed us about locomotion in a way similar to femoral trabecular traits if another individual or more individuals were chosen. Namely, SB would have been distinct compared to all the other taxa, through one single PC_TP+BP_ that is driven by low DA (similarly to femoral data) and by high BV/TV. The latter would represent the opposite pattern compared to the femoral one (i.e., low BV/TV in SB; see above). However, considering the strong loads experienced by the forelimbs in suspensory taxa— even in primates, where this may reverse the typical ‘hindlimb dominance’ characteristic of the group (Granatosky, 2016)—the humeral pattern could also be interpreted in functional terms (also considering that proximal humeral BV/TV has no relationship with body mass proxy, see Results). In this framework, all subfossil lemurs would occupy the extreme peripheral range of the SB morphospace region. It would occur especially for *Palaepropithecus* (both *P. ingens* and *P. kelyus*) while *Mesopropithecus* and *Babakotia* would occupy less distinctive regions. If more extreme positions in the SB region correspond to more extreme suspensory/bridging adaptations, then the humeral data results for subfossil lemurs would be totally consistent with previous locomotor reconstruction (Marchi *et al*. 2016 and references). Likewise, the more extreme position of *M. madagascariensis* compared to *M. edwarsi* would suggest more extreme suspensory/bridging arboreal habits in the former — small body-sized koala-lemur — compared to the latter —large body-sized koala-lemur. However, this would partially contradict what we proposed based on femoral trabecular structure (see above). Future work investigating already available or newly unearthed postcranial material of subfossil lemurs is needed to clarify these puzzling patterns. It may be time to consider that the exceptional features of subfossil lemurs — such as their unusually large body mass relative to strepsirrhine standards — may make locomotor inference based solely on extant analogues limiting, as has been suggested for other peculiar mammalian groups (e.g. xenarthrans, Vizcaíno *et al*. 2018)

Finally, it should be briefly discussed the single observations representing exceptions to the main patterns for femoral trabecular structure that we found (Figs. 5-7 and above). For instance, the chimpanzee *Pan troglodytes,* in this work categorized as QSC, often lies within the range interpreted as characterizing SB for femoral head (i.e. higher PC2_TP+BP_ scores), lateral condyle (i.e. higher PC3_TP+BP_ scores) and medial condyle (i.e. higher PC3_TP+BP_ scores). The chimpanzee shows some suspensory and bridging in its locomotor repertoire (Table S3, Granatosky, 2018; Hunt, 2022, and references) and it explains its position in the locomotor behavior PCA, i.e. tending more, compared to *Gorilla gorilla*, to high PC1 scores, which higher portions are occupied by SB, Fig. 1A). Also, the black howler *Alouatta caraya* (AG category, in this work) and the eastern woolly lemur *Avahi laniger* (L category, in this work) lie within the SB range of variation for proximal femoral and medial condyle data, respectively (Figs. 5, 7), the latter also mirrored by *Gorilla gorilla* (QSC category, in this work). All these taxa, although assigned to other locomotor categories, rarely exhibit either bridging (*A. caraya* and *A. laniger*) or suspensory behavior (e.g., *Gorilla gorilla*), but not both, as is the case for *P. troglodytes* (Table S3) that, accordingly, lies within SB range more often (see above). These patterns, on the one hand support the relationship between femoral trabecular properties and suspensory/bridging locomotion, on the other hand highlight again the common issue of locomotor discrete categories. In this work we designed categories based on distribution on PCA biplot in turn deriving from locomotor quantitative data (Fig 1A), since we ultimately needed discrete categories for the classification technique that we used, i.e. pDFA. However, assigning taxa to a single group inevitably obscures aspects of their locomotor diversity—for example, the suspensory and/or bridging behaviors of African great apes. The latter are particularly difficult to categorize in broad mammalian contexts, as they display distinctive and nearly unique locomotor adaptations—such as knuckle-walking (Tarrega-Saunders *et al*. 2021)—that resist classification into broad discrete categories. The complexity related to African great ape locomotor categorization is also reflected by the fact that their category in this work, i.e. QSC, often occupies wide morphospace regions, diagnostic of broad intra-category variability. Future works should aim to include quantitative behavioral data and analyze them as such, directly analyzing the covariation between morphological and locomotor matrices.

## Conclusions

In this work, we analyzed the humeral and femoral trabecular structure of sloth- and koala-lemurs, addressing this aspect of their anatomy for the first time in a morphofuncitonal and comparative context, using a broad sample of extant mammals. We found that femoral trabecular structure distinguishes arboreal mammals that predominantly employ suspensory and bridging behaviors—such as tree sloths, lorisids, and orangutans. These taxa exhibit substantially more isotropic trabeculae and lower bone volume fraction, traits that can be functionally interpreted as adaptations to their specialized arboreal habits.

Accordingly, we found that the largest palaeopropithecid, *Palaeopropithecus*, was strongly adapted to a suspensory/bridging arboreal repertoire, aligning with previous reconstructions. However, incomplete or missing data prevented confirmation of previously reconstructed behaviors for other palaeopropithecids, namely *Babakotia* and *Mesopropithecus*. As for koala-lemurs, we could not provide a detailed reconstruction for the small body-sized *Megaladapis madagascariensis*, here only inferred as non-suspensory/bridging. Strikingly, however, we found that the largest species, *Megaladapis edwardsi*, was a specialized suspensory and bridging arboreal mammal. More studies are needed, but nevertheless, our results provide evidence for one of the largest mammals ever known to show adaptations to an arboreal lifestyle characterized by the use of suspensory and bridging behavior—potentially matched only by extant orangutans. These new insights substantiate the use of trabecular bone structure to clarify past locomotor habits and highlight the importance of studying or recovering additional postcranial elements from extinct primates.

## Author contributions

Fabio Alfieri (collection, processing, and analysis of μCT data, performing statistical analyses, drafting, writing and editing of the manuscript, conceiving the study, designing the sample, planning statistical analysis steps, interpreting results), Julia Arias-Martorell, Carla Argilés-Esturgó and Damiano Marchi (conceiving the study, designing the sample, planning statistical analysis steps, interpreting results, contributing to writing and editing of the manuscript).

## Ackowledgements

For allowing visits to museum collections, CT-scanning, providing and/or allowing the download of μCT data we thank: Matthew Skinner, Timothy Ryan (at the time of data acquisition funded by the NSF grant number BCS-0617097), Heiko Temming, the AMNH (New York) Mammalogy Department and Joshua Wisor, Frieder Mayer, Christiane Funk, Anna Rosemann, Kristin Mahlow, Martin Kirchner (MfN, Berlin, Germany), Eva Bärmann and Jan Decher (ZFMK, Bonn, Germany), Frank Zachos and Alexander Bibl (NMW, Vienna, Austria), Guillaume Billet (MNHN, Paris, France), Renaud Lebrun (MRI-ISEM, Montpellier, France), Neil Duncan and Sara Ketelsen (AMNH, New York, USA), Adam Ferguson, William Simpson (FMNH), April Isch Neander and Zhe-Xi Luo (Univ. of Chicago) (Chicago, USA), Matt Borths, Catherine Riddle (DPC) and Justin Gladman (SMIF) (Durham, NC, USA), Rachel Jennings (PC-MER, Birchington-on-Sea, UK), Linda Gordon (Smithsonian NMNH). We additionally thank the Bavarian State Collection of Zoology (ZSM, Munich, Germany)—in particular Anneke H. van Heteren and Gerhard Haszprunar—for access to the objects and the Department of Human Evolution, Max Planck Institute for Evolutionary Anthropology, Leipzig, Germany (Jean-Jacques Hublin) and the Tai Chimpanzee Project (Christophe Boesch). Moreover, we thank the MRI platform member of the national infrastructure France-BioImaging supported by the French National Research Agency (ANR-10-INBS-04, “Investment for the future”), the labex CEMEB (ANR-10-LABX-0004) and NUMEV (ANR-10-LABX-0020). This work was performed in part at the Duke University Shared Materials Instrumentation Facility (SMIF), a member of the North Carolina Research Triangle Nanotechnology Network (RTNN), which is supported by the National Science Foundation (award number ECCS-2025064) as part of the National Nanotechnology Coordinated Infrastructure (NNCI). During part of this work, F.A. was part of the Department of Earth Sciences (University of Cambridge, Cambridge, UK).

## Data availability statement

Raw results and additional information on the studied specimens (Tables S1-S2, including MorphoSource ARK identified to download CTscan data), quantitative locomotor data on the studied species (Table S3), summary statistics for all the PCAs (Tables S4-S14) and additional methods and results (Supporting Information) can be downloaded from https://figshare.com/s/ab30e10e8ebd7c0e7b31

## Supporting Information

Supporting Information (Appendices S1-S10, Tables S15-S19): additional methods and additional results

– Table S1-S2: additional information and raw results for humeral (Table S1) and femoral (Table S2) data
– Table S3: Quantitative data on locomotor behaviors (taken and processed as detailed in the Main Text and Supporting Information), together with the locomotor categories to which species were assigned.
– Table S4: Summary statistics for the PCA performed on quantitative locomotor data
– Table S5: Summary statistics for the PCA performed on proximal humerus trabecular data
– Table S6: Summary statistics for the PCA performed on distal humerus trabecular data
– Table S7: Summary statistics for the PCA performed on proximal femur trabecular data
– Table S8: Summary statistics for the PCA performed on femoral lateral condyle trabecular data
– Table S9: Summary statistics for the PCA performed on femoral medial condyle trabecular data
– Table S10: Summary statistics for the PCA performed on proximal humerus trabecular data + body mass proxy
– Table S11: Summary statistics for the PCA performed on distal humerus trabecular data + body mass proxy
– Table S12: Summary statistics for the PCA performed on proximal femur trabecular data + body mass proxy
– Table S13: Summary statistics for the PCA performed on femoral lateral condyle trabecular data + body mass proxy
– Table S14: Summary statistics for the PCA performed on femoral medial condyle trabecular data + body mass proxy

## References

Alfieri F. Convergent Evolution of Humeral and Femoral Functional Morphology in Slow Arboreal Mammals. PhD dissertation. Humboldt-Universität zu Berlin, HU, Germany. 2022.

Alfieri F., Botton-Divet L., Nyakatura J.A., Amson E.. Integrative approach uncovers new patterns of ecomorphological convergence in slow arboreal xenarthrans. Journal of Mammalian Evolution 2022; 29: 283–312

Alfieri F., Botton-Divet L., Wölfer J. et al. A macroevolutionary common-garden experiment reveals differentially evolvable bone organization levels in slow arboreal mammals. Communications Biology 2023; 6(**1**): 995.

Alfieri F., Demuth O.E., Steell E.M. et al. Constrained diversification of avian wing bone internal architecture. Functional Ecology 2025b; DOI: 10.1111/1365-2435.70178

Alfieri F., Nyakatura J.A., Amson E. Evolution of bone cortical compactness in slow arboreal mammals. Evolution 2021; 75, 542–554.

Alfieri F., Veneziano A., Panetta D. et al. The relationship between primate distal fibula trabecular architecture and arboreality, phylogeny and size. Journal of Anatomy 2025a; 246(**6**), 907– 935.

Amson, E., Arnold, P., van Heteren, A.H. et al. Trabecular architecture in the forelimb epiphyses of extant xenarthrans (Mammalia). Frontiers in Zoology 2017; 14:52.

Amson E., Nyakatura J.A. The postcranial musculoskeletal system of xenarthrans: insights from over two centuries of research and future directions. Journal of Mammalian Evolution 2018; 25, 459–484.

Arias-Martorell J., Zeininger A., Kivell T.L. Trabecular structure of the elbow reveals divergence in knuckle-walking biomechanical strategies of African apes. Evolution 2021; 75(**11**), 2959– 2971.

Baab K.L., Perry J.M.G., Rohlf F.J., Jungers W.L. Phylogenetic, ecological, and allometric correlates of cranial shape in Malagasy lemuriforms. Evolution 2014; 68:1450–1468.

Barak M.M. Bone modeling or bone remodeling: That is the question. American Journal of Physical Anthropology; 2019 172(**2**): 153–155.

Barak M.M., Lieberman D.E., Hublin J.J. Of mice, rats and men: trabecular bone architecture in mammals scales to body mass with negative allometry. Journal of Structural Biology; 2013 183: 123–131.

Barak M.M., Lieberman D.E., Raichlen D., et al. Trabecular evidence for a human-like gait in *Australopithecus africanus*. PLoS ONE 2013; 8: e77687.

Bicca_Marques J.C., Calegaro_Marques C. Behavioral thermoregulation in a sexually and developmentally dichromatic neotropical primate, the black_and_gold howling monkey (*Alouatta caraya*). American Journal of Physical Anthropology 1998; 106(**4**): 533–546.

Biewener A.A., Fazzalari N.L., Konieczynski D.D., Baudinette R.V. Adaptive changes in trabecular architecture in relation to functional strain patterns and disuse. Bone 1996; 19: 1–8.

Boyer D.M., Toussaint S., Godinot M. Postcrania of the most primitive euprimate and implications for primate origins. Journal of Human Evolution 2017; 111: 202–215.

Boyer D.M., Yapuncich G.S., Butler J.E., et al. Evolution of postural diversity in primates as reflected by the size and shape of the medial tibial facet of the talus: evolution of postural diversity in primates. American Journal of Physical Anthropology 2015; 157: 134–177.

Carlson K.J. Investigating the form-function interface in African apes: Relationships between principal moments of area and positional behaviors in femoral and humeral diaphyses. American Journal of Physical Anthropology 2005; 127: 312–334.

Carlson K.J., Lublinsky S., Judex S. Do different locomotor modes during growth modulate trabecular architecture in the murine hind limb? Integrative and Comparative Biology 2008; 48: 385– 393.

Chang G., Pakin S.K., Schweitzer M.E., et al. Adaptations in trabecular bone microarchitecture in Olympic athletes determined by 7T MRI. Journal of Magnetic Resonance Imaging 2008; 27(**5**): 1089–1095.

Clavel J., Escarguel G., Merceron G. mvmorph: an R package for fitting multivariate evolutionary models to morphometric data. Methods in Ecology and Evolution 2015; 6: 1311–1319.

Cotter M.M., Simpson S.W., Latimer B.M., Hernandez C.J. Trabecular microarchitecture of hominoid thoracic vertebrae. The Anatomical Record 2009; 292: 1098–1106.

Crowley B.E. A refined chronology of prehistoric Madagascar and the demise of the megafauna. Quaternary Science Reviews 2010; 29.19–20: 2591–2603.

Demes B., Carlson K.J., Franz T.M. Cutting corners: the dynamics of turning behaviors in two primate species. Journal of Experimental Biology 2006; 209(5): 927–937.

Demes B., Jungers W.L., Wunderlich R.E., et al. Body size and leaping kinematics in Malagasy vertical clingers and leapers. Journal of Human Evolution 1996; 31: 367e388.

Demes B., Larson S.G., Stern Jr. J.T., et al. The kinetics of primate quadrupedalism: “hindlimb drive” reconsidered. Journal of Human Evolution 1994; 26: 353e374.

Domander R., Felder A.A., Doube M. BoneJ2 - refactoring established research software. Wellcome Open Research 2021; 6: 37.

Doube M., Kłosowski M.M., Wiktorowicz-Conroy A.M., et al. Trabecular bone scales allometrically in mammals and birds. Proceedings of the Royal Society B: Biological Sciences 2011; 278: 3067–3073.

Douglass K., Hixon S., Wright H.T., et al. A critical review of radiocarbon dates clarifies the human settlement of Madagascar. Quaternary Science Reviews 2019; 221: 105878.

Dunn R.H. Functional Morphology of the Postcranial Skeleton. In: Methods in Paleoecology. Reconstructing Cenozoic Terrestrial Environments and Ecological Communities.

Croft D.A., Su D.F., Simpson S.W. (eds), Springer, Cham, 2018: 23–36.

Fajardo R.J., DeSilva J.M., Manoharan R.K., et al. Lumbar vertebral body bone microstructural scaling in small to medium_sized strepsirhines. The Anatomical Record 2013 296(2): 210– 226.

Fajardo R.J., Müller R. Three-dimensional analysis of nonhuman primate trabecular architecture using micro-computed tomography: Primate Trabecular Bone and the 3-D μCT Method. American Journal of Physical Anthropology 2001; 115: 327–336.

Fajardo R.J., Müller R., Ketcham R.A., Colbert M. Nonhuman anthropoid primate femoral neck trabecular architecture and its relationship to locomotor mode. The Anatomical Record 2007; 290: 422–436.

Fleagle J.G. Primate Adaptation and Evolution - 3rd Edition. Academic press, 2013.

Furnell S., Blanchard M.L., Crompton R.H., Sellers W.I. Locomotor ecology of *Propithecus verreauxi* in Kirindy Mitea National Park. Folia Primatologica 2015; 86(4): 223–230.

Gebo D.L. Vertical clinging and leaping revisited: Vertical support use as the ancestral condition of strepsirrhine primates. American Journal of Physical Anthropology 2011; 146: 323–335.

Gebo D.L. Locomotor diversity in prosimian primates. American Journal of Physical Anthropology 1987; 13: 271–281.

Gebo D.L. The Anatomy of the Prosimian Foot and its Application to the Primate Fossil Record. Ph.D. dissertation. Duke University, 1986.

Georgiou L., Dunmore C.J., Bardo A. et al. Evidence for habitual climbing in a Pleistocene hominin in South Africa. The Proceedings of the National Academy of Sciences (PNAS) 2020; 117(15): 8416–8423.

Georgiou L., Kivell T.L., Pahr D.H., et al. Trabecular architecture of the great ape and human femoral head. Journal of Anatomy 2019; 234: 679–693.

Georgiou L., Kivell T.L., Pahr D.H., Skinner M.M. Trabecular bone patterning in the hominoid distal femur. PeerJ 2018; 6: e5156.

Godfrey L.R. Adaptive diversification of Malagasy strepsirrhines. Journal of Human Evolution 1988; 17. 1–2: 93–134.

Godfrey L.R., Granatosky M.C., Jungers W.L. The Hands of Subfossil Lemurs. In: The Evolution of the Primate Hand. Anatomical, Developmental, Functional, and Paleontological Evidence. Kivell T.L., Lemelin P., Richmond B.G., Schmitt D (eds). Springer, New York, 2016: 421– 453.

Godfrey L.R., Jungers W.L. The extinct sloth lemurs of Madagascar. Evolutionary Anthropology 2003; 12: 252–263.

Godfrey L.R., Jungers W.L. Quaternary fossil lemurs. In: The Primate Fossil Record. Hartwig W.C. (ed) Cambridge University Press, Cambridge, 2002: 97–121.

Godfrey L.R., Jungers W.L., Burney, D.A. Subfossil Lemurs of Madagascar. In: Cenozoic Mammals of Africa. Werdelin L. and Sanders W.J. (eds). Univ of California Press, 2010: 351–367.

Godfrey L.R., Jungers W.L., Reed K.E. et al. Subfossil Lemurs: Inferences About Past and Present Primate Communities. In: Natural Change and Human Impact in Madagascar. Goodman, S.M., Patterson, B.D. (Eds.). Smithsonian Institution Press, Washington, DC, 1997: 218–256.

Godfrey L.R., Sutherland M.R., Paine R.R., et al. Limb joint surface areas and their ratios in Malagasy lemurs and other mammals. American Journal of Physical Anthropology 1995; 97 (1): 11–36.

Gommery D., Ramanivosoa B., Tombomiadana-Raveloson S., et al. Une nouvelle espèce de lémurien géant subfossile du Nord-Ouest de Madagascar (*Palaeopropithecus kelyus*, Primates). Comptes Rendus Palevol 2009, 8: 471–480.

Gônet J., Laurin M., Hutchinson J.R. Evolution of posture in amniotes–Diving into the trabecular architecture of the femoral head. Journal of Evolutionary Biology 2023; 36(8): 1150–1165.

Granatosky M.C. Pedal Morphology and Locomotor Behavior of the Subfossil Lemurs of Madagascar. In: The Evolution of the Primate Foot: Anatomy, Function, and Palaeontological Evidence. Zeininger A., Hatala, K. G., Wunderlich, R. E., & Schmitt, D (eds), Cham: Springer International Publishing, 2022: 415–440.

Granatosky M.C. Primate Locomotion. In: Encyclopedia of Animal Cognition and Behavior. J. Vonk & T. Shackelford (eds), Cham: Springer International Publishing, 2020: 1–7.

Granatosky M.C. A review of locomotor diversity in mammals with analyses exploring the influence of substrate use, body mass and intermembral index in primates. Journal of Zoology 2018; 306: 207–216.

Granatosky M.C. A Mechanical Analysis of Suspensory Locomotion in Primates and Other Mammals. Doctoral dissertation. Duke University. NC. USA, 2016.

Granatosky M.C., Miller C.E., Boyer D.M., Schmitt D. Lumbar vertebral morphology of flying, gliding, and suspensory mammals: Implications for the locomotor behavior of the subfossil lemurs *Palaeopropithecus* and *Babakotia*. Journal of Human Evolution 2014; 75: 40–52.

Hamrick M.W., Simons E.L., Jungers W.L. New wrist bones of Malagasy giant subfossil lemurs. Journal of Human Evolution 2000; 38: 635–650.

Hanna J.B., Granatosky M.C., Rana P., Schmitt D. The evolution of vertical climbing in primates: evidence from reaction forces. The Journal of Experimental Biology 2017; 220: 3039–3052.

Harrigan T.P., Mann R.W. Characterization of microstructural anisotropy in orthotropic materials using a second rank tensor. Journal of Materials Science 1984; 19: 761–767.

Herrera J.P., Dávalos L.M. Phylogeny and divergence times of lemurs inferred with recent and ancient fossils in the tree. Systematic Biology 2016; 65: 772–791.

Hogg R.T., Godfrey L.R., Schwartz G.T., et al. Lemur biorhythms and life history evolution. PLoS ONE 2015; 10 (8): e0134210.

Huiskes R., Ruimerman R., van Lenthe G.H., Janssen J.D. Effects of mechanical forces on maintenance and adaptation of form in trabecular bone. Nature 2000; 405: 704–706.

Hunt K.D. Primate Suspensory Behavior. In: Encyclopedia of Animal Cognition and Behavior. Cham: Springer International Publishing, 2022: 5598–5604.

Ishida H., Hirasaki E., Matano S. Locomotion of the Slow Loris Between Discontinuous Substrates. In: Topics in Primatology. Matano, S., Tuttle, R. H., Ishida, H. & Goodman, M. (eds), University of Tokyo Press., Tokyo, 1992: 139–152.

Ishida H., Jouffroy F., Nakano Y. Comparative Dynamics of Pronograde and Upside Down Horizontal Quadrupedalism in the Slow Loris (Nycticebus coucang). In: Gravity, Posture and Locomotion in Primates. Jouffroy F., Stack M,, Niemitz C. (eds), Firenze, Il Sedicesimo, 1990: 209–220.

Jungers W.L. Adaptive diversity in subfossil Malagasy prosimians. Zeitschrift für Morphologie und Anthropologie 1980; 71: 177–186.

Jungers W.L. The functional significance of skeletal allometry in *Megaladapis* in comparison to living prosimians. American Journal of Physical Anthropology 1978; 49: 303–314.

Jungers W.L. Hindlimb and pelvic adaptations to vertical climbing and clinging in *Megaladapis*, a giant subfossil prosimian from Madagascar. Yearbook of Physical Anthropology 1977; 20: 508–524.

Jungers W.L., Demes B., Godfrey L.R. How Big Were the “Giant” Extinct Lemurs of Madagascar? In: Elwyn Simons: A Search for Origins. J. G. Fleagle & C. C. Gilbert (eds.), Springer-Verlag, 2008: 343–360.

Jungers W.L., Godfrey L.F., Simons E.L., et al. Phylogenetic and functional affinities of *Babakotia* (Primates), a fossil lemur from northern Madagascar. The Proceedings of the National Academy of Sciences (PNAS) 1991; 88: 9082–9086.

Jungers W.L., Godfrey L.R., Simons E.L., Chatrath P.S. Phalangeal curvature and positional behavior in extinct sloth lemurs (Primates, Palaeopropithecidae). The Proceedings of the National Academy of Sciences (PNAS) 1997; 94: 11998–12001.

Jungers W.L., Godfrey L.R., Simons E.L. et al. Ecomorphology and Behavior of Giant Extinct Lemurs from Madagascar. In: Reconstructing Behavior in the Primate Fossil Record. Plavcan J.M., Kay R.F., Jungers W.L., van Schaik C.P. (eds), Kluwer Academic/Plenum Publishers, New York, 2002: 371–411.

Jungers W.L., Lemelin P., Godfrey L.R., et al. The hands and feet of *Archaeolemur*: metrical affinities and their functional significance. Journal of Human Evolution 2005; 49.1: 36–55.

Karanth K.P., Delefosse T., Rakotosamimanana B., et al. Ancient DNA from giant extinct lemurs confirms single origin of Malagasy primates. The Proceedings of the National Academy of Sciences (PNAS) 2005; 102: 5090–5095.

Keaveny T.M., Morgan E.F., Niebur G.L., Yeh O.C. Biomechanics of trabecular bone. Annual Review of Biomedical Engineering 2001; 3: 307–333.

Khosla S., Melton J.L., Achenbach S.J., et al. Hormonal and biochemical determinants of trabecular microstructure at the ultradistal radius in women and men. The Journal of Clinical Endocrinology & Metabolism 2006; 91: 885–891.

Kimura T. Hindlimb dominance during primate high-speed locomotion. Primates 1992; 33: 465e476.

Kimura T. Bipedal and Quadrupedal Walking of Primates: Comparative Dynamics. In: Primate Morphophysiology, Locomotor Analyses and Human Bipedalism. Kondo, S. (ed.). University of Tokyo Press, Tokyo, 1985: 81–104.

Kimura T., Okada M., Ishida H. Kinesiological Characteristics of Primate Walking: its Significance in Human Walking. In: Environment, Behavior and Morphology: Dynamic Interactions in Primates. Morbeck, M.E., Preuschoft, H., Gomberg, N. (eds.), G. Fischer, New York, 1979: 297–311.

Kistler L., Ratan A., Godfrey L.R., et al. Comparative and population mitogenomic analyses of Madagascar’s extinct, giant ‘subfossil’ lemurs. Journal of Human Evolution 2015; 79: 45–54.

Kivell T.L. A review of trabecular bone functional adaptation: what have we learned from trabecular analyses in extant hominoids and what can we apply to fossils? Journal of Anatomy 2016; 228: 569–594.

Kivell T.L., Davenport R., Hublin J.-J., et al. Trabecular architecture and joint loading of the proximal humerus in extant hominoids, *Ateles*, and *Australopithecus africanus*. American Journal of Physical Anthropology 2018; 167: 348–365.

Klingenberg C.P. Methods for studying allometry in geometric morphometrics: a comparison of performance. Evolutionary Ecology 2022; 36.4: 439–470.

Larson S.G. Unique Aspects of Quadrupedal Locomotion in Nonhuman Primates. In: Primate Locomotion. Strasser, E., Fleagle, J., Rosenberger, A., McHenry, H. (eds.). Plenum Press, New York, 1998: 157–173.

Lieberman D.E., Devlin M.J., Pearson O.M. Articular area responses to mechanical loading: effects of exercise, age, and skeletal location American Journal of Physical Anthropology 2001; 116(4): 266–277.

Macho G.A., Abel R.L., Schutkowski H. Age changes in bone microstructure: do they occur uniformly? International Journal of Osteoarchaeology 2005; 15: 421–430.

MacLatchy L., Müller R. A comparison of the femoral head and neck trabecular architecture of *Galago* and *Perodicticus* using micro-computed tomography (μCT). Journal of Human Evolution 2002; 43: 89–105.

Maquer G., Musy S.N., Wandel J., et al. Bone volume fraction and fabric anisotropy are better determinants of trabecular bone stiffness than other morphological variables. Journal of Bone and Mineral Research 2015; 30: 1000–1008.

Marchi D., Ruff C.B., Capobianco A., et al. The locomotion of *Babakotia radofilai* inferred from epiphyseal and diaphyseal morphology of the humerus and femur. Journal of Morphology 2016; 277: 1199–1218.

Marciniak S., Mughal M.R., Godfrey L.R. et al. Evolutionary and phylogenetic insights from a nuclear genome sequence of the extinct, giant, “subfossil” koala lemur Megaladapis edwardsi. The Proceedings of the National Academy of Sciences (PNAS) 2021; 118: e2022117118.

Marshall S.K., Spainhower K.B., Sinn B.T., et al. Hind limb bone proportions reveal unexpected morphofunctional diversification in xenarthrans Journal of Mammalian Evolution 2021; 28(3): 599–619.

Martin R.D. Origins, diversity and relationships of lemurs. International Journal of Primatology 2000; 21: 1021–1049.

Mielke M., Wölfer J., Arnold P., et al. Trabecular architecture in the sciuromorph femoral head: allometry and functional adaptation. Zoological Letters 2018; 4: 10.

Modlesky C.M., Subramanian P., Miller F. Underdeveloped trabecular bone microarchitecture is detected in children with cerebral palsy using high-resolution magnetic resonance imaging. Osteoporosis International 2023; 19: 169–176.

Monclús-Gonzalo O., Alba D.M., Duhamel A., et al. Early euprimates already had a diverse locomotor repertoire: Evidence from ankle bone morphology. Journal of Human Evolution 2023; 181: 103395.

Monclús-Gonzalo O., Alba D. M., Fabre A. C., Marigó J. Reconstruction of the locomotor repertoire of early primates in the light of astragalar and calcaneal shape. Journal of Human Evolution 2010; 206: 103730.

Muldoon K.M. Paleoenvironment of Ankilitelo Cave (late Holocene, southwestern Madagascar): implications for the extinction of giant lemurs. Journal of Human Evolution 2010; 58.4: 338– 352.

Muldoon K.M., Godfrey L.R., Jungers W.L., Chipman J.W. Geographic patterning in subfossil primate community dynamics in Madagascar. American Journal of Physical Anthropology 2009; S48: 195.

Nowak R.M. Walker’s Mammals of the World (v.1). Baltimore: Johns Hopkins University Press, 1999.

Nyakatura J.A. The convergent evolution of suspensory posture and locomotion in tree sloths. Journal of Mammalian Evolution 2012; 19: 225–234.

Nyakatura J.A., Alfieri F., Granatosky M.C. et al. Upside-Down and in Slow Motion: The Functional Morphology and Mechanics of Sloth Suspensory Locomotion. In: Functional Morphology and Biomechanics of Arboreal Locomotion in Tetrapods. M.C Granatosky, S. Toussaint and D. Youlatos (eds). Springer Nature Editions. In press. pre-print: DOI: 10.13140/RG.2.2.35257.89449

Orlando L., Calvignac S., Schnebelen C., et al. DNA from extinct giant lemurs links archaeolemurids to extant indriids. BMC Evolutionary Biology 2008; 8: 1–9.

Patel B.A., Goodenberger K.E., Boyer D.M., Jungers W.L. Hallucal reduction in sloth lemurs and morphological convergence on orangutans by *Palaopropithecus*. American Journal of Physical Anthropology 2013; 56(Suppl)::217.

Pearson O.M., Lieberman D.E. The aging of Wolff’s “law”: ontogeny and responses to mechanical loading in cortical bone. American Journal of Physical Anthropology 2004; 125(S39): 63–99.

Pinheiro J., Bates D., DebRoy S., Sarkar D. nlme: Linear and Nonlinear Mixed Effects Models. R package version 3.1–147. https://rdrr.io/cran/nlme/, 2020.

Plasse M., Amson E., Bardin J., et al. Trabecular architecture in the humeral metaphyses of non_avian reptiles (Crocodylia, Squamata and Testudines): lifestyle, allometry and phylogeny. Journal of Morphology 2019; 280: 982–998.

Polly P.D., Lawing A.M., Fabre A.C., Goswami A. Phylogenetic Principal Components Analysis and Geometric Morphometrics. Hystrix, the Italian Journal of Mammalogy 2013; 24(1): 33–41.

Pontzer H., Lieberman D.E., Momin E. et al. Trabecular bone in the bird knee responds with high sensitivity to changes in load orientation. The Journal of Experimental Biology 2006; 209: 57–65.

Prates H.M., Hass G.P., Bicca-Marques J.C. Ranging Behavior of Black-And-Gold Howler Monkeys (Alouatta caraya) in Anthropogenic Habitat Patch in Southern Brazil. In: La Primatología En Latinoamérica 2 – A Primatologia Na America Latina 2. Tomo I Argentina-Colombia.

Ediciones IVIC. Urbani B, Kowalewski M, Cunha RGT, de la Torre S & L Cortés-Ortiz (eds.), Instituto Venezolano de Investigaciones Científicas (IVIC). Caracas, Venezuela, 2018: 259–266.

Qiao Z., Zhou L., Huang J.Z. Sparse linear discriminant analysis with applications to high dimensional low sample size data. International Journal of Applied Mathematics 2009; 39: 48e60.

R Core Team. R: A language and environment for statistical computing. R Foundation for Statistical Computing, Vienna, Austria. https://www.R-project.org/. 2023

Rafferty K.L., Ruff C.B. Articular structure and function in *Hylobates*, *Colobus*, and *Papio*. American Journal of Physical Anthropology 1994; 94: 395–408.

Rayadin Y., Spehar S.N. Body mass of wild Bornean orangutans living in human_dominated landscapes: Implications for understanding their ecology and conservation. American Journal of Physical Anthropology 2015 157(2): 339–346.

Revell L.J.. phytools: an R package for phylogenetic comparative biology (and other things). Methods in Ecology and Evolution 2012; 3: 217–223.

Rolvien T., Hahn M., Siebert U., et al. Vertebral bone microarchitecture and osteocyte characteristics of three toothed whale species with varying diving behaviour. Scientific Reports 2017; 7(1): 1604.

Rook L., Bondioli L., Köhler M., et al. *Oreopithecus* was a bipedal ape after all: evidence from the iliac cancellous architecture. The Proceedings of the National Academy of Sciences (PNAS) 1999; 96(15): 8795–8799.

Rowe N. The Pictorial Guide to the Living Primates. Pogonias Press, East Hampton, New York, 1996.

Rueden C.T., Schindelin J., Hiner M.C., et al. ImageJ2: ImageJ for the next generation of scientific image data. BMC Bioinformatics 2017; 18(1): 529.

Ruff C. Hindlimb articular surface allometry in hominoidea and *Macaca*, with comparisons to diaphyseal scaling. Journal of Human Evolution 1988; 17: 687–714.

Runestad J.A. Postcranial adaptations for climbing in Loridae (Primates). Journal of Zoology 1997; 242: 261–290.

Ryan T.M., Carlson K.J., Gordon A.D. et al. Human-like hip joint loading in *Australopithecus africanus* and *Paranthropus robustus*. Journal of Human Evolution 2018; 121: 12–24.

Ryan T.M., Ketcham R.A. Angular orientation of trabecular bone in the femoral head and its relationship to hip joint loads in leaping primates. Journal of Morphology 2005; 265: 249– 263.

Ryan T.M., Ketcham R.A. Femoral head trabecular bone structure in two omomyid primates. Journal of Human Evolution 2002a; 43: 241–263.

Ryan T.M., Ketcham R.A. The three-dimensional structure of trabecular bone in the femoral head of strepsirrhine primates. Journal of Human Evolution 2002b; 43: 1–26.

Ryan T.M., Krovitz G.E. Trabecular bone ontogeny in the human proximal femur. Journal of Human Evolution 2006 51: 591–602.

Ryan T.M., Raichlen D.A., Gosman J.H. Structural and Mechanical Changes in Trabecular Bone During Early Development in the Human Femur and Humerus. In: Bones: Bone Formation and Development in Anthropology. Percival, C. J., Richtsmeier, J. T. Building (eds). Cambridge: Cambridge University Press 2017: 281–302.

Ryan T.M., Shaw C.N. Gracility of the modern *Homo sapiens* skeleton is the result of decreased biomechanical loading. Proceedings of the National Academy of Sciences (PNAS) 2015; 112: 372–377.

Ryan T.M., Shaw C.N. Trabecular bone microstructure scales allometrically in the primate humerus and femur. Proceedings of the Royal Society B: Biological Sciences 2013; 280: 20130172.

Ryan T.M., Shaw C.N. Unique suites of trabecular bone features characterize locomotor behavior in human and non-human anthropoid primates. PLoS ONE 2012; 7: e41037.

Ryan T.M., Walker A. Trabecular bone structure in the humeral and femoral heads of anthropoid primates. The Anatomical Record 2010; 293: 719–729.

Saers J.P.P., Cazorla-Bak, Y., Shaw, C.N., et al. Trabecular bone structural variation throughout the human lower limb. Journal of Human Evolution 2016; 97: 97–108.

Saparin P., Scherf H., Hublin J.J., et al. Structural adaptation of trabecular bone revealed by position resolved analysis of proximal femora of different primates. The Anatomical Record 2011; 294: 55e67.

Scherf H., Harvati K., Hublin J.-J. A comparison of proximal humeral cancellous bone of great apes and humans. Journal of Human Evolution 2013; 65: 29–38.

Schmitt D., Hanna J.B. Substrate alters forelimb to hindlimb peak force ratios in primates. Journal of Human Evolution 2004; 46: 239e254.

Shapiro L.J., Jungers W.L., Godfrey, L.R., Simons, E.L. Vertebral morphology of extinct lemurs. American Journal of Physical Anthropology 1994; 37: Suppl:179–180.

Shapiro L.J., Seiffert C.V.M., Godfrey L.R., et al. Morphometric analysis of lumbar vertebrae in extinct Malagasy strepsirrhines. American Journal of Physical Anthropology 2005; 128: 823– 839.

Shaw C.N., Ryan T.M. Does skeletal anatomy reflect adaptation to locomotor patterns? Cortical and trabecular architecture in human and nonhuman anthropoids. American Journal of Physical Anthropology 2012; 147: 187–200.

Simons E.L., Godfrey L.R., Jungers W.L., et al. A new giant subfossil lemur, *Babakotia*, and the evolution of the sloth lemurs. Folia Primatologica 1992; 58: 197–203.

Skedros J.G., Baucom S.L. Mathematical analysis of trabecular “trajectories” in apparent trajectorial structures: The unfortunate historical emphasis on the human proximal femur. Journal of Theoretical Biology 2007; 244: 15e45.

Smith R., Jungers W. Body mass in comparative primatology. Journal of Human Evolution 1997; 32: 523–559.

Sokal R., Rohlf F. *Biometry: The Principles and Practice of Statistics in Biological Research* 3rd ed., Freeman, New York, 1995.

Somers K.M. Allometry, isometry and shape in principal components analysis. Systematic Zoology 1989; 38.2: 169–173.

Spoor F., Garland T., Krovitz G. et al. The primate semicircular canal system and locomotion. The Proceedings of the National Academy of Sciences (PNAS) 2007; 104: 10808–10812.

Stauber M., Rapillard L., van Lenthe G.H., et al. Importance of individual rods and plates in the assessment of bone quality and their contribution to bone stiffness. Journal of Bone and Mineral Research 2006; 21(4): 586–595.

Szalay F.S., Delson E. Evolutionary History of the Primates. New York: Academic Press, 1979.

Tarrega-Saunders E.L., King C., Roberts A.M., Thorpe S.K. Knuckle-walking and behavioural flexibility in great apes. Revue de Primatologie 2021; 12.

Tattersall I. The Primates of Madagascar. Columbia University Press, New York, 1982.

Tattersall I. Notes on the Cranial Anatomy of the Subfossil Malagasy Lemurs. In: Lemur Biology. I Tattersall, and RW Sussman (eds.). New York: Plenum Press, 1975: 111–124.

Tattersall I. Subfossil lemuroids and the “adaptive radiation” of the Malagasy lemurs. Transactions of the New York Academy of Sciences 1973; 35: 314–324.

Tattersall I., Schwartz J.H. Craniodental morphology and systematics of the Malagasy lemurs (Primates, Prosimii). Anthropological Papers of the American Museum of Natural History 1974; 52: 137–192.

Thorpe S.K., Crompton R.H. Locomotor ecology of wild orangutans (Pongo pygmaeus abelii) in the Gunung Leuser Ecosystem, Sumatra, Indonesia: A multivariate analysis using log_linear modelling. American Journal of Physical Anthropology 2005; 127(1): 58–78.

Tsegai Z.J., Kivell T.L., Gross T., et al. Trabecular bone structure correlates with hand posture and use in hominoids. PLoS ONE 2013; 8: e78781.

Tsegai Z. J., Skinner M.M., Pahr D.H. et al. Systemic patterns of trabecular bone across the human and chimpanzee skeleton. Journal of Anatomy 2018; 232(4): 641–656.

Tsegai Z. J., Skinner M.M., Pahr D.H. et al. Ontogeny and variability of trabecular bone in the chimpanzee humerus, femur and tibia. American Journal of Physical Anthropology 2018; 167: 713–736.

Upham N.S., Esselstyn J.A., Jetz W. Inferring the mammal tree: species-level sets of phylogenies for questions in ecology, evolution, and conservation. PLoS Biology 2019; 17: e3000494.

Vizcaíno S.F., Toledo N., Bargo M.S. Advantages and limitations in the use of extant xenarthrans (Mammalia) as morphological models for paleobiological reconstruction. Journal of Mammalian Evolution 2018; 25: 495–505.

Volpato V., Viola T.B., Nakatsukasa M., et al. Textural characteristics of the iliac-femoral trabecular pattern in a bipedally trained Japanese macaque. Primates 2008; 49: 16–25.

Vuillame-Randriamanantena M., Godfrey L.R., Jungers W.L., Simons E.L. Morphology, taxonomy and distribution of *Megaladapis*-giant subfossil lemur from Madagascar. Comptes Rendus de l’Académie des Sciences Série III 1992; 315: 1835–1842.

Walker A. Prosimian Locomotor Behavior. In: The Study of Prosimian Behavior. Doyle GA. and Martin RD. (eds), Academic Press, Inc, 1979: 543–565.

Walker A., 1974. Locomotor Adaptations in Past and Present Prosimian Primates. In: Primate Locomotion. Jenkins F.A. (ed). Academic Press, New York: 349–381.

Walker A. Locomotor Adaptations in Recent and Fossil Madagascan Lemurs. PhD. thesis, University of London, 1967

Walker A., Ryan T.M., Silcox M.T., et al. The semicircular canal system and locomotion: The case of extinct lemuroids and lorisoids. Evolutionary Anthropology 2008; 17: 135–145.

Wunderlich R.E., Jungers W.L., Godfrey L.R., Simons E.L. Pedal form and function in subfossil indroids. American Journal of Physical Anthropology. 1994; Suppl. 18:211.

Wunderlich R.E., Simons E.L., Jungers W.L. New pedal remains of *Megaladapis* and their functional significance. American Journal of Physical Anthropology 1996; 100: 115–139.

Yapuncich G.S., Granatosky M.C. Footloose: Articular surface morphology and joint movement potential in the ankles of lorisids and cheirogaleids. American Journal of Physical Anthropology 2021; 175(4): 876–894.

Yoder A.D., Rakotosamimanana B., Parsons T.J. Ancient DNA In Subfossil Lemurs. In: New Directions in Lemur Studies. B. Rakotosamimanana, H. Rasamimanana, J. U. Ganzhorn, & S. M. Goodman (eds.). Springer: 1999:1–17.

Youlatos D., Pylarinos D., Karantanis N.E., Rychlik L. Locomotion, postures, and substrate use in captive southern pygmy slow lorises (Strepsirrhini, Primates): implications for conservation. Animals 2025; 15(11): 1576.

Young J.W. Ontogeny of limb force distribution in squirrel monkeys (*Saimiri boliviensis*): insights into the mechanical bases of primate hind limb dominance. Journal of Human Evolution 2012; 62.4: 473–485.

Zihlman A.L., Mcfarland R.K., Underwood C.E. Functional anatomy and adaptation of male gorillas (*Gorilla gorilla gorilla*) with comparison to male orangutans (*Pongo pygmaeus*). The Anatomical Record 2011; 294(11): 1842–1855.

